# Sonic Hedgehog Is An Important Regulator Of Intervertebral Disc Homeostasis And Rejuvenation

**DOI:** 10.64898/2026.09.18.752770

**Authors:** Sarthak Mohanty, Rajakumar Anbazhagan, Paul Pricop, Sarah Loh, Robert Pinelli, Tom Ross, Eric A. Bogner, James C. Farmer, Russel C. Huang, Darren R. Lebl, Bernard A. Rawlins, Han Jo Kim, Harvinder S. Sandhu, Matthew E. Cunningham, Sheeraz Qureshi, Todd J. Albert, Chitra L. Dahia

**Affiliations:** Orthopedic Soft Tissue Research Program, Hospital for Special Surgery, New York City, NY, USA, 10021; Spine Service, Hospital for Special Surgery, New York City, NY, USA, 10021; Weill Cornell Medical College, New York City, NY, USA, 10021; Department of Cell and Developmental Biology, Weill Cornell Medical College, New York City, NY, USA, 10021

**Keywords:** sonic hedgehog, brachyury, nucleus pulposus, intervertebral disc degeneration, regeneration, annulus fibrosus, innervation

## Abstract

Intervertebral disc degeneration and associated neurological symptoms constitute a global health burden, yet no cure is currently available ^1,2^. Each disc consists of a central nucleus pulposus, surrounded by annulus fibrosus, and end plates connecting it to the growth plates of adjacent vertebral bodies. The notochord-descendant nucleus pulposus continues to express sonic hedgehog, which regulates the proliferation and differentiation of all components of neonatal mouse discs ^3–7^. Sonic hedgehog expression and function decline with age and are associated with disc pathologies, including terminal differentiation of nucleus pulposus cells to chondrocyte-like phenotype ^7,8^. Here, we used fate-mapping and conditional genetic mouse models to test the role of sonic hedgehog in the homeostasis of aging discs. We found that loss of NP-derived *Shh* is sufficient to accelerate multiple features of age-associated disc degeneration, including reduced NP cell number, altered matrix turnover, and induction of inflammatory, angiogenic and neurotrophic programs. Age-related disc pathologies were more prevalent in the lumbosacral discs of mice, like in humans ^8–13^. Also, like mice, the expression of sonic hedgehog and its targets by human nucleus pulposus cells declines with age and pathological degeneration. Moreover, pharmacologic Hedgehog activation partially restored anabolic gene expression and reduced catabolic, inflammatory and neurotrophic mediators in degenerated human NP explants ex vivo. These findings indicate that Sonic hedgehog, a developmental signal retained in the adult NP, is functionally active and required for disc homeostasis during aging and supports activation of hedgehog signaling as a candidate disease-modifying pathway for intervertebral disc pathologies.

## INTRODUCTION

Aging is a significant risk factor for various chronic degenerative and inflammatory diseases, including those of the musculoskeletal system, which are becoming more prevalent and impacting the quality of life of our increasing aged population ^1,2^. However, little is known about the cellular and molecular basis of age-related musculoskeletal tissue pathologies, including those affecting intervertebral discs of the spine, a leading cause of chronic back pain. Intervertebral discs (“discs” hereafter) form a cartilaginous joint in the spine, providing flexibility to the spine while resisting compressive forces during movement and maintaining space for the entry and exit of peripheral nerve fibers. The central region of each disc is occupied by the proteoglycan-rich nucleus pulposus (NP), surrounded by orthogonal layers of collagenous annulus fibrosus (AF), which are together sandwiched between cartilaginous end plate (EP) that connects the disc to the growth plates of the vertebral body on its proximal and distal ends (**Fig. 1a** and **c**). The extracellular matrix (ECM) is an essential component of disc structure and function ^14,15^. Pathologies of the lumbar disc, including hypocellularity, loss of ECM, and narrowing of disc height, increase with age and are often associated with chronic back pain. The disc comprises the largest avascular, aneural tissue in the body; but the AF of degenerated human discs become vascularized and innervated along with increased expression of neurotropic and inflammatory factors ^16–21^. AF of aged mouse disc also becomes vascularized and innervated ^12^. The root cause of these degenerative pathologies is primarily unknown.

**Figure 1.**
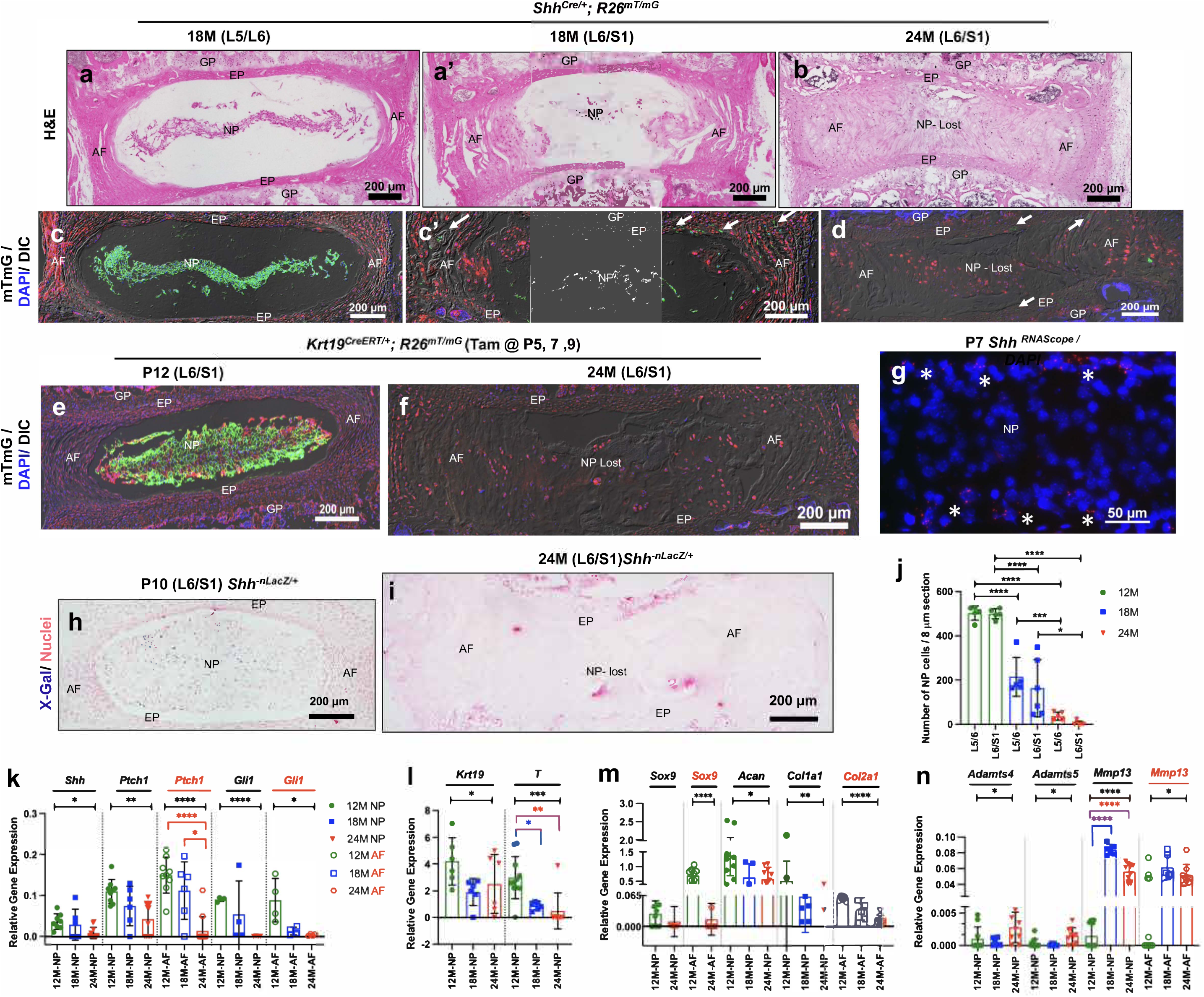
Disc degeneration is associated with loss of *Shh* expression. Representative images from each cohort are imaged in the mid-coronal plane. Lumbosacral discs of 18-month-old (18M, **a**, **a’**, **c**, **c’**) and 24-month-old (24M, **b**, **d**) *Shh^Cre/+;^ R26^mT/mG^* mice (*n* = 3 per age cohort). The mid-coronal sections from 18M (**a** and **a’**) and 24M old (**b**) *Shh^Cre/+;^ R26^mT/mG^* mice were stained with H&E to visualize histological changes. Serial sections were stained with DAPI and fate-mapped mGFP+ NP cells and changes in disc structure (**c**, **c’**, and **d**) were visualized at 20X magnification using epifluorescence and dark-field (DIC) imaging. Mid-coronal sections of the lumbosacral (L6/S1) disc of P12 (**e**) and 24M (**f**) old *Krt19^CreERT/+;^ R26^mT/mG^* mice that were induced with tamoxifen at P5 to mark NP cells as mGPF+ for fate-mapping (*n* = 3 per age cohort) and imaged at 20X magnification. RNA *in situ* for *Shh* (RNAscope) at P7 (**g**) and imaged at 60X magnification; asterisks (*) mark a sub-set of *Shh-* expressing NP cells (TOM+). Nuclei are counterstained with DAPI (blue) in **c**, **c’**, **d**, **e**, **f**, and **g**. LacZ staining of lumbar discs from P10 (**h**) and 24M old mice (**i**) *Shh*-*^nLacZ^* reporter mice imaged at 20X magnification show *Shh*-expressing cells (blue). Nuclei are counterstained with Nuclear Fast Red in **i** and **j**. Quantification of the number of NP cells in lumbosacral discs from 12M, 18M, and 24M old mice (**j**). Multiplex qPCR results of NP and AF cells microdissected from lumbar discs from 12M, 18M, and 24M old FVB mice using TaqMan probes for genes indicated at top (**k** - **n**) (*n* = 6 to 11 mice per age cohort) and *B2m* as internal control. Each dot represents a biological replicate in **j** - **n**. One-way ANOVA followed by Tukey’s multiple comparisons test (**j**). Brown-Forsythe and Welch ANOVA tests followed by Dunnett’s T3 multiple comparisons test (**k** - **n**). The black horizontal line and black Asterisks (*) above the bars indicate ANOVA results, while the colored line and Asterisks shows the results from multiple comparisons between cohorts (**k** - **n**). Data are presented as mean ± S.D. \**P* < .05; ** *P* < .01; *** *P* < .001; **** *P* < .0001. NP, nucleus pulposus. AF, annulus fibrosus. EP, end plate. GP, growth plate.

The role of key developmental signaling pathways, including hedgehog (Hh), Wnt, TGFb, BMP, and FGF in regulating patterning and growth of the soft musculoskeletal tissues inducing differentiation of intervertebral disc ^4,6,22,23^, growth plate (GP) (^24^, reviewed by ^25^), and tendon ^26–29^ is well established. Developmental signaling pathways that pattern musculoskeletal tissues may also serve adult homeostatic functions, but whether their age-associated decline causally contributes to disc degeneration remains unresolved. Identifying molecular regulators that maintain adult disc homeostasis may reveal disease-modifying strategies for age-associated disc degeneration.

NP cells of the discs are descendants of the embryonic notochord ^30^. Notochord is an important signaling center and via SHH expression regulates patterning during early embryogenesis (reviewed by ^31^). Postnatal NP cells continue to express SHH ^5,6^ and regulate all components of the disc, including NP, AF, and EP, shown by expression and regulation of its downstream targets PTCH1 and GLI1 in these tissues ^3,4,7^. SHH regulates NP cell proliferation and differentiation through ECM production by NP, AF, and EP cells in lumbar discs of neonatal mice ^4,7^ and sacral discs of skeletally mature mice ^3^. Expression of *Shh* declines with age and is associated with the differentiation of reticular-shaped NP cells into chondrocyte-like cells (CLCs) isolated in lacunae by 16-18 months (M) of age ^7,8^. However, the fate of NP cells in naturally aging and pathological discs and the role of SHH in aging discs remain unknown. Here, using independent lineage-tracing and conditional genetic approaches, we tested whether NP-derived SHH functions as an adult homeostatic signal whose decline contributes causally to age-associated disc degeneration, and whether Hh pathway activation can preserve or partially restore features of disc homeostasis in mouse and human tissue.

## RESULTS

### Decline in *Shh* expression is associated with disc pathologies in aging mice

Humans have five lumbar (L) discs, while most strains of mice have six. Disc pathology is prevalent in the lumbosacral discs of human [L3 to sacral level 1 (S1)] ^9–11^ as well as in mice the (L5 to S1) ^8,9,12,32,33^, hence we focused on L5-S1 discs for fate-mapping and histological analysis in mice. Using lineage-tracing strategies, we determined the fate of NP cells in the lumbar discs of aged mice. We first employed *Shh^Cre/+^*; *R26^mT/mG^* alleles where the recombination is expected to occur when *Shh* is first expressed in the node (notochord precursor) at E7.5 ^34^, permanently marking all *Shh*-expressing cells as mGFP+ that can be traced over time in the NP of mouse discs (^30^, **Extended data Fig.1a**). While H&E staining of an 18-month-old mouse disc showed normal histology of L5/6 disc with numerous NP cells, the adjacent L6/S1 disc showed fewer NP cells (**Fig. 1a** and **a’**), along with modest invasion of AF lamella into NP space (**Fig. 1a’**). However, by 24 months of age, the disc was filled with lamellar tissue, and no distinct NP space or cells were observed in the center of the disc (**Fig. 1b**). At 18-months of age epifluorescence microscopy along with dark-field imaging showed that all NP cells were reticular-shaped and homogenously mGFP+ with organized AF layers in the L5/6 discs (**Fig. 1c, Extended data Fig. 1b, e, h**, and **k**). The adjacent L6/S1 disc of the same mouse had fewer and smaller mGFP+ NP cells, and AF exhibited disintegrated lamella (**Fig. 1c’, Extended data Fig. 1b’, e’, h’**, and **k’**). At 24 months, all mGFP+ cells were lost in the lumbosacral discs (**Fig. 1d, Extended data Fig. 1c, f, I**, and **l**). Using lineage-tracing in combination with immunostaining for cytokeratin 19 (KRT19), a hallmark marker of NP cells ^4,8,35,36^, we observed that all NP cells were mGFP+ (**Fig. 1c, c’**) and continued to express KRT19 with age (**Extended data Fig. 1b, b’**). We did not observe any cells in the NP region that were not descendants of *Shh*-expressing cells or were mGFP-negative and TOM+ (**Fig. 1e, e’, Extended data Fig. 1b, b’**), indicating that in the mice discs, the *nuclei pulposi* is exclusively composed of the notochord-descendant NP cells. Within the limits of our lineage-tracing approach, we found no evidence for substantial replacement of the NP compartment by non-NP-lineage cells before advanced structural collapse. KRT19 immunoreactivity persisted in morphologically intact NP-lineage cells, supporting its use as a marker of maintained NP identity during aging. In naturally aged discs, GFP+ anuclear debris were observed within the NP space and near EP/AF regions, suggesting impaired removal of NP cellular remnants during advanced disc aging (white arrows in **Fig. 1c’** and **d**), indicating an age-related impairment in excretion and clearing of NP cellular content from the disc.

Next, we employed an orthogonal genetic approach to lineage-trace NP cells using *Krt19^CreERT/+^*^;^ *R26^mT/mG^* alleles. In these mice, tamoxifen-induction at P5 permanently turned “on” mGFP expression in the NP cells as shown at P12 (**Fig. 1e**), allowing lineage-tracing of NP cells over time ^8,36^. Tamoxifen-treated *Krt19^CreERT/+^*^;^ *R26^mT/mG^* littermates were allowed to age naturally, and lumbar discs were analyzed at 24 months. Epifluorescence and dark-field imaging showed loss of mGFP+ NP-lineage cells in severely affected lumbosacral discs, supporting depletion rather than fibrotic transdifferentiation of NP cells during advanced aging and degeneration (**Fig. 1f**).

The expression of *Shh* mRNA declines from E12.5 in notochord to P0 in NP cells ^37^ and continues to further decline from P4 to middle-age in mice NP cells ^7,8^. As the relative expression of *Shh* continues to decline, we hypothesized that the dramatic decline in *Shh* expression is due to a lower number of *Shh*-expressing NP cells in postnatal stage. We performed RNA *in situ* analysis at P7 and observed that only a subset of NP cells (∼23%) expressed *Shh* (white asterisk, **Fig. 1g**). Next, using an *Shh-^nLacZ/+^* reporter allele ^38^ that allows visualization of *Shh*-expressing cells, we validated that *Shh*-expression was restricted to a subset of NP cells. X-gal staining showed that ∼20% (±2.01) of NP cells expressed *Shh* (blue) at P10 (**Fig. 1h**), which further declined by 12 months (**Extended Data Fig. 2a**) and was absent in the L6/S1 disc of 24-month-old mice, a point at which the disc was fused (**Fig. 1i**). In addition, using tamoxifen-inducible *Shh^CreERT2/+^; R26^mT/mG^* alleles, we assessed *Shh*-expressing NP cells at 12, 18, and 24 months of age and found that only a subset of NP cells was positive for *Shh-* expression at 12 and 18 months (∼12%, **Extended Data Fig. 2c - e**). This *Shh*-positive subset of NP cells was either lost by 24 months of age (**Extended Data Fig. 2f**) or clumped, forming a syncytium in the center of the disc (**Extended Data Fig. 2f-l’**). However, no mGFP+ cells were observed in the AF and EP regions of the disc at any age analyzed, indicating that the mGFP+ cells observed in **Fig. 1c’-d** and **Extended data Fig. 1** using *Shh^Cre/+^*; *R26^mT/mG^* alleles were NP cells or their debris migrating into AF and EP regions with age. Moreover, before the complete fusion of the disc, we observed *Shh*-expressing, as well as KRT19-IF positive NP cells clumped in the center of the disc (**Extended data Fig. 2g, i, i’**), providing evidence that the age-related disc fusion is due to loss of NP cells along with invasion by the surrounding cells in the space. Quantification of the total numbers of NP cells from 12 to 24 months of age, in all genetic cohorts studied here, revealed a significant loss from 12 to 18 months of age (*P* < .0001) at both L5/L6 and L6/S1 levels and with most NP cells lost by 24 months of age (**Fig.1j**). Our findings show that loss of *Shh*-expressing NP cells is associated with a significant increase in the histopathological score among all components of the disc (**Extended data Fig. 1n** and **n’**). Together, these results suggest that the lamellar tissue occupying the collapsed NP region in advanced degeneration is unlikely to derive from NP-lineage cells (**Fig. 1b, d, f, I, Extended data Fig.1c, f, i**, and **l**), as reported in the injury model of mouse accelerated disc degeneration^39^.

Although neonatal and middle-aged NP cells are morphologically similar and share a history of *Shh*-lineage expression, we observed that only a subset retains detectable *Shh* expression, revealing molecular heterogeneity within the NP compartment. Previous studies have reported decreased *Shh* mRNA expression between neonatal and middle-aged mouse NP cells ^7,8^. Therefore, we tested the mRNA expression of *Shh* and its downstream targets in NP cells from middle-aged (12 months), early aging (18 months), and aged (24 months) mice. NP cells were collected from the proximal lumbar discs as they have more NP cells even in aged mice ^8,9,12^. Multiplex qPCR analysis showed that the loss of *Shh*-expressing NP cells was associated with a decline in *Shh* mRNA expression from 12 to 24 months of age (*P* = .0187, Fig. 1k). Moreover, the decline in *Shh* expression was associated with a similar decrease in mRNA expression of its downstream targets *Ptch1 (*NP, *P = .0032*; AF, *P <.0001)* and *Gli1* (NP, *P* < .0001; AF, *P* = .0492) by NP and AF cells from 12 to 24 months of age (**Fig. 1k**) indicating that Hh signaling in these cells declines with age. Next, we analyzed the expression of known SHH targets KRT19 and Brachyury (*T, Bra, or Tbxt* in humans) in NP cells as they are only expressed by NP cells ^4–6^. We observed a significant decline in mRNA expression of *Krt19* (*P* = .0330) and *T* (*P* = .0009) from 12 to 24 months of age (**Fig. 1l**). Furthermore, immunoreactivity for KRT19, which is expressed by all NP cells at all ages, was detectable until mGFP+ lineage-traced NP cells were present in the discs (18 months in this experiment) but was undetectable (*P* = .0008) coincident with the loss of mGFP+ NP cells in 24-month-old mouse discs (**Extended data Fig. 1b-d**). This result validated the qPCR findings and established KRT19 as robust marker to identify NP cells even in aging and pathological discs.

ECM is a significant component of healthy discs. SHH is reported to regulate the expression of ECM markers in young mouse discs ^3,4^. Here we tested the expression of ECM markers in the discs of aging mice as SHH expression declines. First, we analyzed the expression of crucial ECM markers in our NP lineage-traced mouse discs. Immunostaining for aggrecan (ACAN), chondroitin sulfate proteoglycan (CSPG), and type X collagen α-1 (COLXA1) showed a significant declines in expression of these ECM markers in all components of the discs, which was associated with the loss of lineage-traced NP cells in the lumbar discs of *Shh^Cre/+^*; *R26^mT/mG^* (**Extended Data Fig. 1e-m**) and *Shh^CreERT2/+^*; *R26^mT/mG^* (**Extended Data Fig. 2j - l’**) mice. Next, the decline in expression of ECM-related proteins was validated at the transcriptional level through multiplex qPCR analysis of NP and AF cells, where the decline in *Shh* expression and signaling was validated (**Fig. 1k**). SOX9 is an upstream regulator of *Col2a1* ^40^ and a downstream target of SHH in NP cells of young mouse discs ^4^. Multiplex qPCR analysis at 12 to 24 months of age showed significant declines in mRNA expression of *Sox9* in AF cells (*P* < .0001)*, Acan* in NP cells *(P* = .0239), *Col1a1* in NP cells (*P* = .0062), and *Col2a1* in AF cells (*P* < .0001, **Fig. 1m**). However, we did not observe a significant change in the expression of *Sox9* in NP cells from 12 to 24 months of age (**Fig. 1m**). In NP cells, reduced expression of ECM markers was associated with significant increases in mRNA expression of aggrecanases including *Adamts4* (*P* = .0420) and *Adamts5* (*P* = .0102, **Fig. 1n**), as well as increases in collagenase *Mmp13* by NP (*P* < .0001) and AF cells *(P* = .0152, **Fig. 1n**), indicating that age-related loss of *Shh*-expressing NP cells affects matrix remodeling in the disc cells. Multiple comparison analyses revealed that the main effects were between 12 and 24 months of age. Together, these results link disc aging to loss of *Shh*-expressing NP cells, reduced Hh pathway activity, diminished NP identity markers and altered matrix turnover, supporting a role for SHH in adult disc homeostasis.

### During natural aging, degenerating discs become innervated and are inflamed

Previous studies reported increased expression of neurotrophic factors, including brain-derived neurotrophic factor (BDNF), nerve growth factor (NGF), substance-P (Tac2), and inflammatory cytokines, including interleukins IL1b and IL6, tumor necrosis factor-alpha (TNFa), and cyclooxygenase-2 (Cox2 of PGE2) in the degenerated disc tissue of patients ^41–44^. Furthermore, neovascularization and innervation are reported in degenerated human discs ^16–21^. Increased expression of these neurotrophic, angiogenic and inflammatory molecules associated with neoinnervation and neovascularization is reported in preclinical animal models of disc degeneration ^12,45,46^. To test whether age-related decline in *Shh* expression is associated with changes in disc homeostasis leading to its innervation and vascularization we employed multiplex qPCR analysis to check the expression of known neurotropic, angiogenic (**Fig. 2a**), and inflammatory markers (**Fig. 2b** and **c**) in the NP and AF cells collected from lumbar disc of 12-, 18- and 24-month-old FVB mice, where a decline in *Shh* expression and response was confirmed (**Fig. 1k**). Results of qPCR analysis showed age-related increased mRNA expression of neurotropic factors including *Bdnf* (NP, *P* = .0226; AF, *P =* .0055), *Ngf* (NP, *P =* .0097), and *Tac2* (NP, *P* = .0058) (**Fig. 2a**). We observed increased mRNA expression of angiogenic factor *Vegfa* by NP cells (*P =* .0033), but not in AF cells (**Fig. 2a**). Furthermore, an age-related increase in mRNA expression of *Cox2* was observed in both NP (*P* = .0138) and AF (*P* = .0022) cells (**Fig. 2b**), indicating an age-related increased inflammatory response in the disc. Multiple comparison analyses revealed that the main effects occurred between 12 and 24 months of age for the neurotropic and inflammatory markers; therefore, we only compared these two age cohorts for subsequent analysis. Next, we tested the expression of key inflammatory cytokines in NP and AF cells, where a decline in *Shh* expression and signaling was validated (**Fig. 1k**). We observed a significant increase in mRNA expression of *Il1b* by NP (*P* < .0001) and AF (*P* = .0337) cells, and *Tnfa* by NP (*P =* .0475) cells but not in AF cells (*P* = .5) from 12 to 24 months of age (**Fig. 2c**). These results indicate that age-associated decline in NP-derived SHH coincides with induction of neurotrophic, angiogenic and inflammatory transcriptional programs in the degenerating disc.

**Figure 2.**
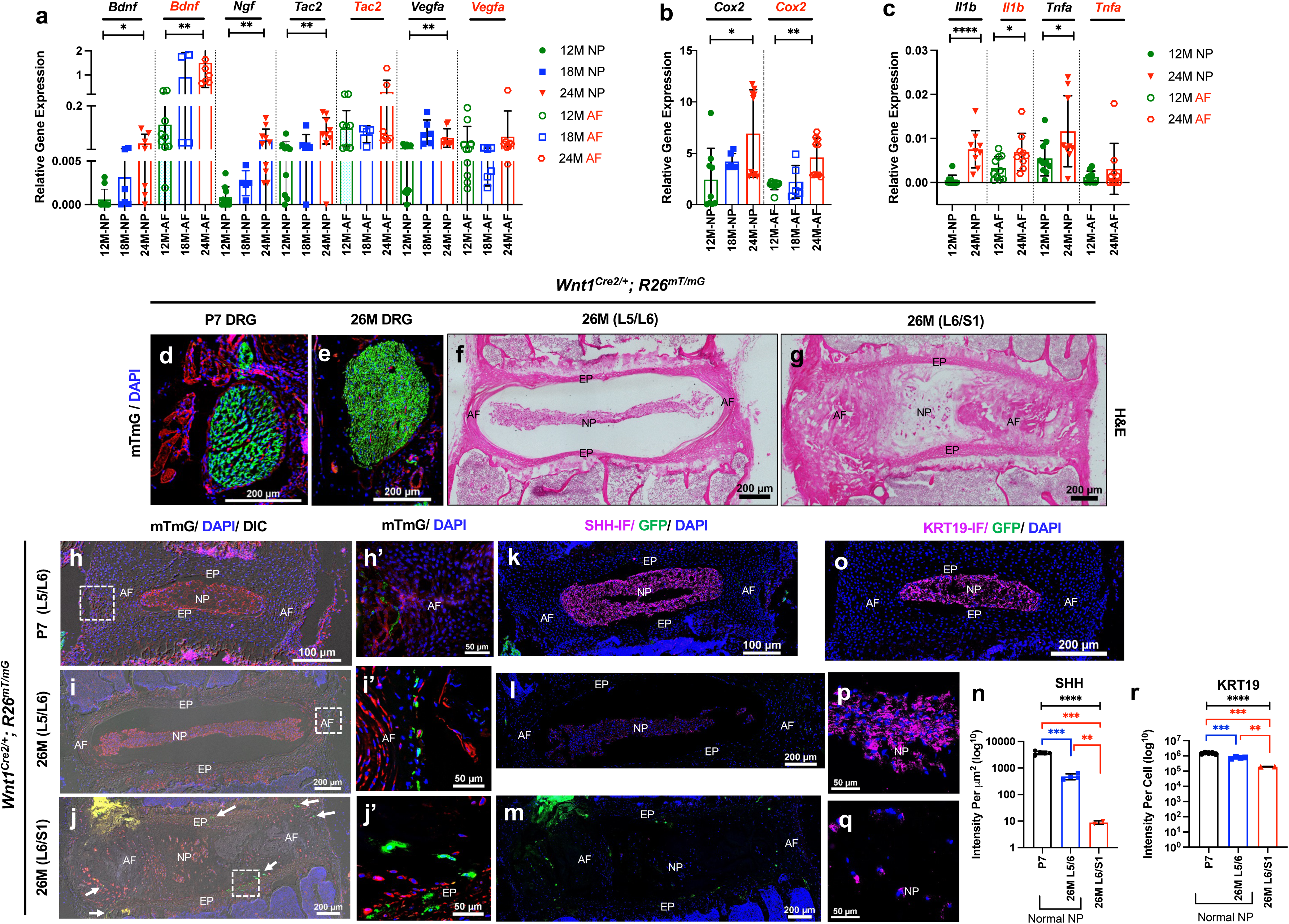
Disc innervation is associated with loss of *Shh* expression and NP cells during aging. Multiplex qPCR results of NP and AF cells collected from lumbar discs of 12-month (12M), 18M, and 24M old FVB mice using TaqMan probes for the genes indicated at the top of the graph (**a - c**) (*n* = 6 to 11 mice per age cohort) and *B2m* as an internal control. Coronal section of lumbosacral DRG from P7 (**d**) and 26M (**e**) old *Wnt1^Cre2/+^; R26^mT/mG^* mice (*n* = 3 per age cohort) showing that all nerve fibers are mGFP+. Representative H&E-stained mid-coronal section of L5/L6 (**f**) and L6/S1 (**g**) disc of 26M old *Wnt1^Cre2/+^; R26^mT/mG^* mice. Mid-coronal section of lumbosacral discs of P7 (**h**, **h’**) and L5/6 (**i**, **i’**) and L6/S1 (**j**, **j’**) of 26M old *Wnt1^Cre2/+^; R26^mT/mG^* mice at 20x (**h**, **I**, **j**) and 60X (**h’**, **i’**, **j’**) magnification. Immunoreactivity for SHH (purple) on the mid-coronal section of lumbosacral discs of P7 (**k**), and L5/6 (**l**) and L6/S1 (**m**) of 26M old *Wnt1^Cre2/+^; R26^mT/mG^* mice at 20x magnification. KRT19 immunoreactivity of the mid-coronal section of lumbosacral discs of P7 (**o**) at 20x magnification, and L5/6 (**p**), and L6/S1 (**q**) of 26M old *Wnt1^Cre2/+;^ R26^mT/mG^* mice discs that show NP cells at 60x magnification. Nuclei are counterstained with DAPI (blue, **d**, **e**, **h**, **h’**, **i**, **i’**, **j**, **j’**, **k**, **l**, **m**, **o**, **p**, **q**). Quantification of mean fluorescence intensity for SHH (**n**) and fluorescence intensity per cell for KRT19 (**r**) at P7 and 26M of age based on the presence of NP cells. Each dot represents a biological replicate in **a** - **c**, **n** and **r**. Brown-Forsythe and Welch ANOVA tests followed by Dunnett’s T3 multiple comparisons test (**a** - **c**, **n** and **r**). The black line and Asterisks (*) above the bars indicate the ANOVA results (**a** - **c**, **n** and **r**), while the colored line shows the results from multiple comparisons (**n** and **r**). Data are presented as mean ± S.D. * *P* < .05; ** *P* < .01; *** *P* < .001. NP, nucleus pulposus. AF, annulus fibrosus. EP, end plate.

Next, to test whether the decline in *Shh* expression and associated increase in expression of neurotrophic factors in pathologically aging mouse discs is also associated with disc innervation, we generated conditional dual-fluorescent reporter to trace sensory neurons using *Wnt1^Cre2/+^; R26^mT/mG^* alleles. *Wnt1^Cre2/+^* induces recombination of *R26^mT/mG^* allele in the neural crest cells, which gives rise to the dorsal root ganglion (DRG), the sensory hub. Consequently, all sensory nerve fibers can be traced by mGFP expression, as shown in the DRGs of P7 and 26-month-old mice (**Fig. 2d** and **e**). We employed the 26-month-old *Wnt1^Cre2/+^*; *R26^mT/mG^* mice to lineage-trace the nerve fibers in the lumbosacral discs. H&E staining showed normal structure and morphology of the L5/6 discs, but severe histopathology including NP hypocellularity and disorganization and bulging of AF lamellae both outwards and inwards in the L6/S1 discs of the 26-month-old *Wnt1^Cre2/+^; R26^mT/mG^* mice (**Fig. 2f** and **g**). Epifluorescence showed very few mGFP+ nerve fibers in the outer region of AF of lumbar discs of P7 and L5/L6 disc of 26-month-old *Wnt1^Cre2/+^; R26^mT/mG^* mice (**Fig. 2h – i’**). In contrast, the severely degenerated L6/S1 disc of the same 26-month-old mouse contained *Wnt1*-lineage mGFP+ sensory fibers extending through the AF and EP regions into the disc space (white arrows in **Fig. 2j**, and **j’**). Immunostaining for SHH on these discs showed that the discs with increased axonal sprouting had a significant decline in SHH immunoreactivity (*P* < .0001, **Fig. 2k-n**). Immunostaining for KRT19 on serial sections of these discs showed significant loss of KRT19 immunoreactivity (*P* < .0001) along with loss of NP cells (**Fig. 2o-r**). As sensory neuron of DRG cells also expresses *Shh* ^47^, the anuclear mGFP+ structures observed in the lumbosacral discs of 24-month-old *Shh^Cre/+^*; *R26^mT/mG^* mice (**Fig. 1d, Extended data Fig. 1f, i**, and **l**) support neural sprouting in these degenerated discs. These observations indicate that loss of SHH expression and NP-lineage cells is associated with an inflammatory, pro-angiogenic and pro-innervation disc microenvironment during advanced aging.

### Conditional targeting of *Shh* in NP cells accelerates pathological disc degeneration

We next tested whether SHH is required for adult disc homeostasis and whether loss of NP-derived SHH is sufficient to accelerate degenerative features observed during natural aging. We assessed this possibility by conditional targeting of *Shh* in NP cells of middle-aged mice. Tamoxifen-mediated recombination was induced in about 10-month-old *Krt19^CreERT/+^*; *Shh^flx/flx^* (*Shh*-cKO) and *Shh^flx/flx^* (wild-type, WT, control) littermates, and effects were analyzed 2.5 and 5 months later. H&E staining of L6/S1 discs showed typical disc structure in control cohorts at both ages (**Fig. 3a – b’**). In *Shh*-cKO mice, NP cells acquired a multinucleated chondrocyte-like morphology ^8^ by 2.5 months after *Shh*-targeting and were markedly depleted by 5 months, accompanied by encroachment of surrounding lamellar tissue and accelerated disc collapse relative to their WT littermate controls (**Fig. 3a – b’**).

**Figure 3.**
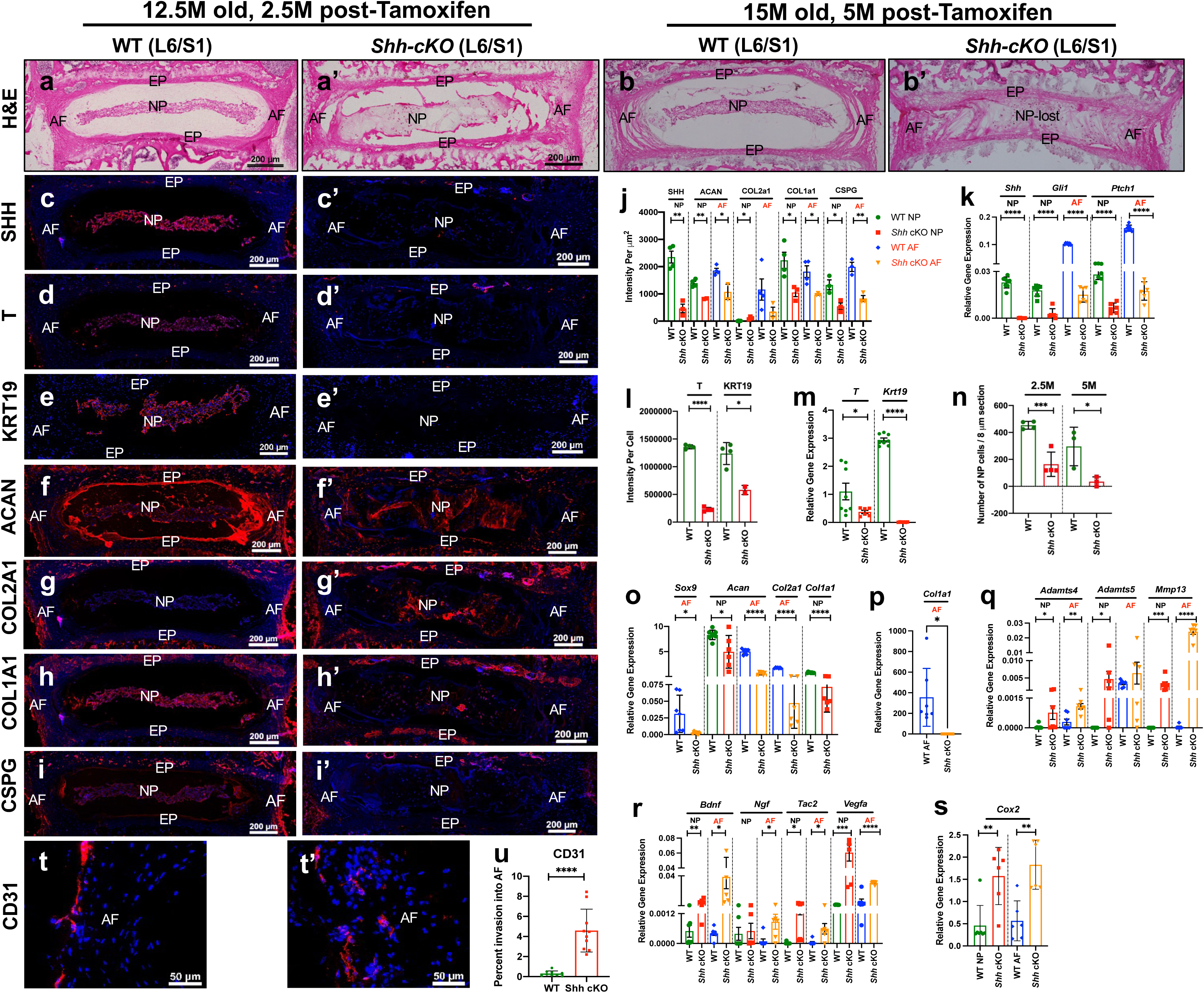
Conditional targeting of *Shh* accelerates disc pathologies. Representative H&E- and immuno-stained (indicated at left) mid-coronal sections of lumbosacral discs (L6/S1) of WT (*Shh^flx/flx^, n* = 7) (**a**, **c** - **i**) and *Shh*-cKO (*Krt19^CreERT/+;^ Shh^flx/flx^, n* = 6) littermates (**a’**, **c’** - **i’**) at 12.5 month (12.5M) of age, and 2.5M post-tamoxifen induction. Representative H&E-stained images captured at 20X magnification show the mid-coronal sections from 15M old mice at 5M post-tamoxifen induction (*n* = 3/cohort, **b**, **b’**). Representative fluorescence images captured at 20X magnification show immunoreactivity (red) for SHH (**c**, **c’**), T (**d**, **d’**), KRT19 (**e**, **e’**), ACAN (**f**, **f’**), COL2A1 (**g**, **g’**), COL1A1 (**h**, **h’**), CSPG (**i**, **i’**) 2.5M post-tamoxifen induction. Representative fluorescence images captured at 60X magnification show immunoreactivity for CD31 in AF regions of L6/S1 disc of WT (**t**) and *Shh*-cKO (**t’**) littermates 2.5M post-tamoxifen induction. Nuclei are counterstained with DAPI (blue, **c – i’, t, t’**). Quantification of immunofluorescence intensity in NP and AF regions (**j**) and per NP cell (**l**) of data presented in panels **c** - **i’**. Quantification of the number of NP cells (**n**). Quantification of percent invasion of CD31-IF+ structure into the AF space (**u**). Multiplex qPCR analysis of NP and AF cells 2.5M post-tamoxifen induction (**k**, **m**, **o**, **p**, **q**, **r, s**) using TaqMan^TM^ probes for genes indicated above the bars, and *B2m* as internal control. Each dot represents a biological replicate in **j** - **s**, **u**. Statistical analysis by unpaired *t-*test. Data presented as mean ± S.D. * *P* < .05; ** *P* < .01; *** *P* < .001; **** *P* < .0001. NP, nucleus pulposus. AF, annulus fibrosus. EP, end plate.

We validated the loss of *Shh* and expression of its targets using the 2.5-month cohort. Efficient targeting and loss of *Shh* in the NP cells of the *Shh*-cKO cohort compared to WT littermates was confirmed by multiplex qPCR analysis of NP cells (*P* < .0001, **Fig. 3k**) and immunostaining (*P* = .0013, **Fig. 3c, c’** and **j**) of the lumbar discs. Multiplex qPCR analysis of Hh signaling targets *Gli1 and Ptch1* showed significant declines in both NP and AF cells of *Shh*-cKO mice compared to WT littermates (all comparisons *P* < .0001, **Fig. 3k**), confirming loss of SHH signaling in these cells following *Shh*-targeting in NP cells. Moreover, a significant decline in the expression of SHH targets in NP cells, including *T* (qPCR, *P* = .05; IF, *P* < .0001) and *Krt19* (qPCR, *P* < .0001; IF, *P* = .012) (**Fig. 3d’, e’, l** and **m**) in *Shh*-cKO mice compared to WT littermates was observed. A significant decline in NP cell number (*P* = .0009 at 2.5 months post-*Shh-*cKO; and *P* = .0372 at 5 months post-*Shh-*cKO) was observed in *Shh-*cKO discs compared to those of age-matched WT littermate (**Fig. 3n**) and was comparable to that observed by 24months of age (**Fig. 1j**). TUNEL assay indicated a significant increase in NP cell death (*P* = .0233) in 2.5 months post-*Shh-*cKO mice compared to WT littermates (**Extended Data Fig. 2**). Most NP cells were already lost by 5 months post-*Shh-*cKO. The disc fusion phenotype in **Fig. 3b’** was like that observed with physiological aging at about 24 months of age (**Fig. 1b, d, f, i’, Extended Data Fig. 1f, i, l**, and **Extended Data Fig. 2f**). These results indicate that residual SHH signaling remains necessary for NP cell survival, maintenance of their identity, and tissue organization in middle-aged discs.

SHH regulates ECM in neonatal mouse discs ^3,4^. ECM is an essential component of the disc that is lost during pathological disc aging along with a decline in *Shh* expression (**Fig. 1k, m, Extended Data Fig. 1e – m**, and **Extended Data Fig. 2i – l’**). Next, we assessed whether decline in ECM turnover with age is a consequence of SHH loss. Immunostaining showed a decline in expression of ACAN (NP, *P* = .0049; AF, *P* = .0216, **Fig. 3f, f’** and **j**), COL2A1 (NP, *P* = .0102, **Fig. 3g, g’** and **j**), COL1A1 (NP, *P* = .0249; AF, *P* = .0211, **Fig. 3h, h’** and **j**), and CSPG (NP, *P* = .0226; AF, *P* = .0044, **Fig. 3i, i’** and **j**) in the lumbar discs of 2.5 months post-*Shh*-cKO cohort compared to WT littermates. Furthermore, qPCR analysis showed a corresponding decline in the transcriptional level of several ECM markers including *Sox9 (*AF, *P* = .0320), *Acan* (NP, *P* = .0158; AF, *P* < .0001), *Col2a1* (AF, *P* < .0001), and *Col1a1* (NP *P* < .0001, **Fig. 3o**, and AF *P* = .0105. **Fig. 3p**) 2.5 months post-*Shh*-cKO cohort compared to WT littermates. Moreover, increased mRNA expression of *Adamts4* (NP, *P* = .0446; AF, *P* = .0027), *Adamts5* (NP, *P* = .0465) and *MMP13* (NP, *P* = .0009; AF, *P* < .0001) was observed following conditional targeting of *Shh* in NP cells (**Fig. 3q**), indicating that SHH continues to regulate matrix remodeling in aged mouse disc. Expression of *Adamts5* in the AF did not change much (**Fig. 3q**). The qPCR analysis also revealed increased mRNA expression of neurotrophic factors *Bdnf* (NP, *P* = .0056; AF, *P* = .021), *Ngf* (AF, *P* = .026), *Tac2* (NP, *P* = .029; AF, *P* = .0174), angiogenic factor *Vegfa* (NP, *P* = .0002; AF, *P* < .0001, **Fig. 3r**) and inflammatory factor *Cox2* (NP, *P* = .0038; AF, *P* = .0014, **Fig. 3s**) in 2.5-month post-*Shh*-cKO mice compared to WT littermate controls. Due to increased expression of *Vegfa*, we immunostained for CD31 endothelial marker and observed a corresponding increase in vascularization of AF following *Shh* knockout in NP cells (*P* < .0001, **Fig. 3t - u**). Additionally, the histopathological scores showed accelerated disc pathologies following *Shh-*cKO in both set of experiments (**Extended Data Fig. 3f** and **g**), and comparable to aged mice (>24 months of age, **Extended Data Fig. 1n**).

Using an independent approach, we validated the role of SHH in middle-aged mouse disc homeostasis along with fate-mapping of *Shh*-expressing NP cells. In *Shh**<u>^Flx^</u>**^/CreERT2^; R26^mT/mG^* (*Shh*-cKO-mGFP+) mice following tamoxifen-mediated recombination at 11 months of age, *Shh* expression was knocked out (*Shh**<u>^Flx^</u>***allele) in the *Shh*-expressing NP cells (*Shh^CreERT2^*allele), along with the activation of mGFP expression (*R26^mT/mG^* allele), which enabled their fate-mapping post-*Shh*-cKO. Tamoxifen-treated *Shh^CreERT2/+^; R26^mT/mG^* littermates served as controls where mGFP was turned “on” by *Shh*-expressing NP cells, but no change in *Shh* expression was induced (*Shh*-WT-mGFP+). The effects of conditionally targeting *Shh* specifically in the *Shh*-expressing sub-population of NP cells were analyzed three and four months later. As shown by epifluorescence and darkfield microscopy, NP cells maintained their reticular shape in the lumbar discs of *Shh*-WT-mGFP+ controls and validate that only a sub-set of NP cells express *Shh* at this age (**Extended Data Fig. 4a - d’**). However, terminal differentiation of NP cells to form a multi-nucleated syncytium (or CLCs) was observed in the lumbar discs of *Shh-* cKO-mGFP+ cells (**Extended Data Fig. 4a - d’**), validating that SHH is necessary for homeostasis of NP cells even in middle-aged mice and that its loss causes terminal differentiation of NP cells into a CLC phenotype. A recent report where conditional knock-down of Hh receptor smoothened in NP cells (using *Krt19^CreERT^*allele) resulted in a similar phenotype ^48^, validating that these effects are a direct consequence of SHH loss.

Together, these results indicate that residual SHH signaling in middle-aged discs is required to maintain disc homeostasis and restrain inflammatory, angiogenic and neurotrophic remodeling. These results also show that pathological disc aging is a direct consequence of SHH loss.

### Transient overexpression of *rShh* in the NP cells rejuvenates the disc and prevents pathological aging

Next, we tested whether transient restoration of NP-derived SHH signaling in middle-aged mice, could rejuvenate the disc and preserve its structure during subsequent aging. We generated a triple transgenic mouse line [*Krt19^CreERT/+^; R26^rtTA/rtTA^; TetO7-rShh* (*rShh*)] to conditionally and transiently overexpress *rShh* (rat *Shh*) in NP cells of middle-aged mice (12 - 13 months of age) for one month following tamoxifen-induction and doxycycline treatment. The effects were analyzed six months later at approximately 19 months of age. Tamoxifen and doxycycline-treated *R26^rtTA/rtTA^; TetO7-rShh* mice served as controls (WT). H&E staining showed that the L6/S1 discs of WT controls continued to naturally age with loss of NP cells, disorganization of AF lamella, and an overall degenerative phenotype (**Fig. 4a**). A one-month pulse of NP-specific rSHH expression preserved lumbosacral disc architecture six months later, including NP cell morphology and AF organization, relative to littermate controls (**Fig. 4a** and **a’**). Lower histopathological scores of *rShh*-rescued discs confirmed preservation of disc structure relative to littermate controls. (**Extended Data Fig. 5**). The immunoreactivity for GFP validates lineage-tracing of NP cells in the *rShh* cohort (**Fig. 4b – b’**). Immunoreactivity for KRT19, a pan-NP marker, showed high and uniform expression by all NP compared to controls indicating healthy state of these cells (*P* = .0053) (**Fig. 4d – e**). Transient *rShh* overexpression was associated with sustained higher expression of SHH at both transcriptional and translational levels (IF, *P* = .0005, qPCR, *P* = .0363; **Fig. 4f, k**, and **l**), along with increased *Ptch1* (*P* = .02) and *Gli1* (*P* = .0056, **Fig. 4l**) expression six months later compared to littermate controls, consistent with durable enhancement of Hh pathway activity. Moreover, following transient overexpression of *rShh* in NP cells, there was elevated expression of molecular markers of NP cells and SHH targets, including *Krt19* (*P* < .0001) and *T* (*P* = .0017), in the *rShh* cohort compared to WT controls (**Fig. 4m**). Analysis of the ECM markers showed higher expression of ACAN (qPCR *P* = .0026, IF *P* = .0064), COL2A1 (qPCR *P* = .0064, IF *P* < .0001), and COL1A1 (qPCR *P* = .0038, IF *P* = .0008) at both transcriptional and translational levels (**Fig. 4g - i’, k** and **n**). However, we observed little change in *Sox9* mRNA (**Fig. 4n**) and CSPG protein expression (**Fig. 4j, j’** and **k**) in rSHH-pulsed discs compared to WT controls. qPCR analysis showed that transient *rShh* overexpression also resulted in decreased mRNA expression of matrix-degrading enzymes like *Adamts4* (*P* = .0016, **Fig. 4o**). We also observed a significant decline in mRNA expression of neurotropic, angiogenic, and inflammatory factors including *Tac2* (*P* = .0106)*, Vegfa* (*P* = .0004)*, IL1b* (*P* = .0112), and *Cox2* (*P* = .0119) in *rShh* discs compared to WT controls (**Fig. 4p** and **q**). The mRNA expression for *Adamts5*, *Mmp13*, *Bdnf* and *Ngf* was not detected by qPCR in the discs of the *rShh* cohort. Hence, one sample *t* and Wilcoxon test was conducted which showed a significant decline in mRNA expression of *Adamts5* (*P* = .0006), *Mmp13* (*P* = .0028), and neurotropic factors *Bdnf* (*P* = .0054), and *Ngf* (*P* <.0001) (**Fig. 4o** and **p**) in rSHH-pulsed discs compared to WT controls. No changes were observed in the expression of *Tnfa* between the discs of the two cohorts (**Fig. 4q**). Moreover, a decline in *Vegfa* mRNA expression was also associated with reduced vascularization analyzed by CD31 immunoreactivity in the AFs of L6/S1 discs from the *rShh* mice compared to WTs, which continued to have age-related vascular invasion (*P* = .0037, **Fig. 4r, r’** and **s**). These results provide genetic proof of principle that enhancing SHH signaling in middle age can preserve features of NP identity, maintain matrix-associated programs, and delay degenerative remodeling over a prolonged period in the entire disc.

**Figure 4.**
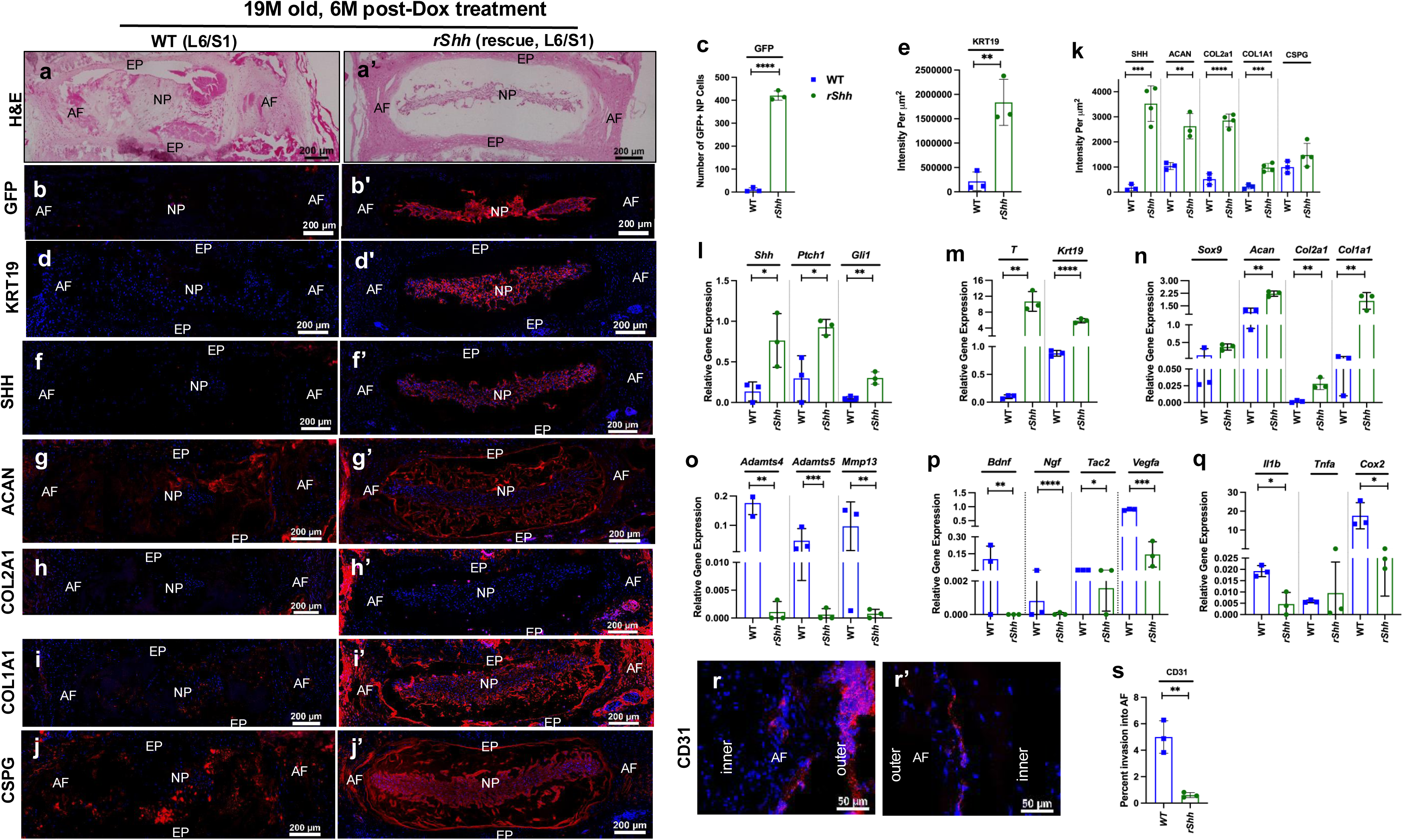
Conditional and transient overexpression of *rShh* prevents age-related disc pathologies. Representative H&E- and immuno-stained (indicated at left) mid-coronal sections of lumbosacral (L6/S1) discs of about 19 month (19M) old WT (*R26^rtTA/rtTA^*; *rShh*, *n* = 3) (**a**, **b**, **d, f** – **j’**); and *rShh* (rescue, *Krt19^CreERT/+^*; *R26^rtTA/rtTA^*; *tetO)_7_CMV-rShh*, *n* = 4) littermates (**a’, b’, d’, f’** - **j’**) six-months post transient over-expression of *rShh* induced by doxycycline treatment. Images captured at 20X magnification. Immunoreactivity (red) for GFP (**b**, **b’**), KRT19 (**d**, **d’**), SHH (**f**, **f’**), ACAN (**g**, **g’**), COL2A1 (**h**, **h’**), COL1A1 (**i**, **i’**), CSPG (**j**, **j’**) imaged at 20X magnification. Immunoreactivity for CD31 in the AF region of L6/S1 disc of WT (**r**) and *rShh* (**r’**) mice captured at 60X magnification on the mid-coronal plane. Nuclei are counterstained with DAPI (blue) in all fluorescence microscopy images. Number of lineage-traced NP cells quantified by GFP immunostaining (**c**). Quantification of immunofluorescence intensity per cell for KRT19 (**e**) and mean immunofluorescence intensity (MFI, **k**) of data presented in figures **f** - **j’**. Multiplex qPCR analysis of lumbar discs of WT (*n* = 3) and *rShh* (*n* = 3) littermates using TaqMan^TM^ probes for genes indicated above horizontal bars and *B2m* as an internal control (**m** - **q**). Quantification of percent invasion of CD31-IF+ structure into the AF space (**s**). Each dot represents a biological replicate in bar graphs. Statistical analysis by unpaired *t-*test with Welch’s correction. qPCR results for *Adamts5*, *Mmp13*, *Bdnf* and *Ngf* were analyzed by one sample *t* and Wilcoxon test. Data are presented as mean ± S.D. * *P* < .05; ** *P* < .01; *** *P* < .001; **** *P* < .0001. NP, nucleus pulposus. AF, annulus fibrosus. EP, end plate.

### *SHH* is crucial for rejuvenating degenerated human NP cells

To test the clinical relevance of SHH expression and signaling, we analyzed NP explants collected as a surgical discard. The pre-surgery MRI images of the lumbar discs were classified using Pfirrmann grades of disc degeneration ^49^. H&E staining showed that increasing Pfirrmann degeneration grade was accompanied by reduced NP cellularity, altered cell morphology and increased CLC phenotype (**Fig. 5a**). NP tissue was collected from males and females 19 - 79 years of age (**Fig. 5b**). Immunostaining (**Fig. 5c**) and multiplex qPCR (**Fig. 5h** and **i**) confirmed *SHH* expression in human NP cells, and its decline with age (*P* = .0486, **Fig. 5h**) and disc degeneration (*P* = .0022, **Fig. 5i**). Kruskal-Wallis test showed that reduction in *SHH* expression was associated with a corresponding decline in the expression of its signaling target *PTCH1* (qPCR, *P* = .0003, **Fig. 5i**) and downstream targets *TBXT (T, or Bra*, qPCR, *P* < .0001, **Fig. 5d** and **j**), *KRT19* (qPCR, *P* = .0003, **Fig. 5j**), ACAN (qPCR, *P* = .0017, **Fig. 5e** and **k**), CSPG (**Fig. 5f**), and COL1A1 (**Fig. 5g**). Moreover, a decline in SHH expression was associated with increased transcriptional levels of *ADAMTS4* (*P* = .0245) and *MMP13* (*P* < .0001, **Fig. 5l**). Higher expression of *BDNF* was previously reported in the AF of human degenerated discs ^20,43,44^. Here, we determined whether higher Pfirrmann degeneration grades were associated with increased expression of neurotrophic factors like mice. qPCR analysis showed increased expression of *BDNF (P* = .0025) and *NGF* (*P* < .0001, **Fig. 5m**) with increased degeneration score of human NP tissue. We also observed increased IL1B expression (*P* = .0003, **Fig. 5n**), consistent with an inflammatory state previously associated with disc degeneration ^20,42–44^.

**Figure 5.**
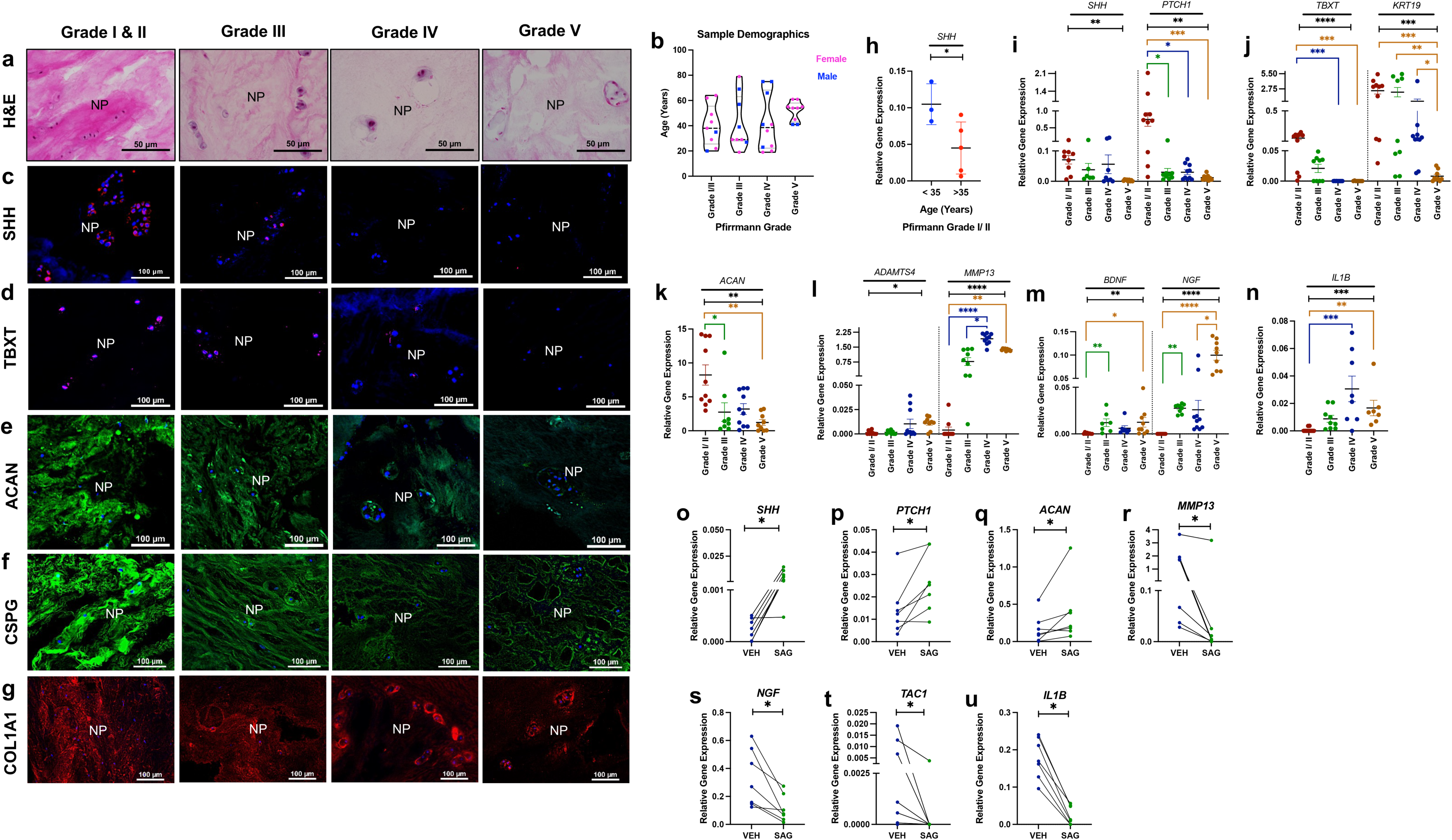
SHH signaling is crucial for the maintenance of human NP cells. H&E (**a**) and immunofluorescence staining (indicated at left, **c** - **g**) of human NP tissues from lumbar discs with Pfirrmann grade I-V (indicated at top) captured at 60x magnification. Sample demographic data showing the distribution of age, gender, and Pfirrmann grade of disc degeneration of NP tissues used in this study (**b**); *n* = 7-9 samples per cohort. Immunoreactivity for SHH (**c**, red), TBXT (**d**, red), ACAN (**e**, green), CSPG (**f**, green), and COL1A1 (**g**, red); nuclei counterstained with DAPI (blue). Multiplex qPCR analysis for *SHH* with *GAPDH* as internal control of NP tissue collected from lumbar discs with Pfirrmann grade I or II (**h**). Multiplex qPCR analysis of NP tissue collected from different disc degenerative stages (**I** - **n**) using TaqMan^TM^ probe for genes indicated above the graphs, and *GAPDH* as internal control. Multiplex qPCR analysis of NP explants cultured for 2.5 days either in the vehicle only (VEH) or stimulated with 5 µM SAG for genes indicated above the graphs (**o** - **u**, *n* = 7) with gene indicated above graphs, and *GAPDH* as internal control. NP tissue from each subject was cultured under two conditions is connected by a line to show pairs. Each dot represents a biological replicate in **b**, **h** - **u**. Statistical analysis by unpaired *t-*test (**h**), Kruskal-Wallis test followed by Dunn’s multiple comparison test (**i** - **n**), Wilcoxon matched-pairs signed rank test (**o** - **u**). The black line and asterisks above the bars indicate ANOVA results; the colored line shows the results from multiple comparisons (**i** - **n**). Data are presented as mean ± S.E.M. * *P* < .05; ** *P* < .01; *** *P* < .001; **** *P* < .0001.

We next tested whether pharmacologic reactivation of Hh signaling could modulate anabolic, catabolic and inflammatory transcriptional programs to rejuvenate the degenerated human NP explants. Because discs with Pfirrmann Grade V had fewer NP cells, NP explants were collected from discs with Pfirrmann Grade III and IV discs. NP tissue collected from each subject was split into two groups; one group was stimulated with a small molecule Hh agonist (SAG, 5 µM) and the other group was treated with equal volume of vehicle as control. Hh signaling activation in human NP explants was evaluated by qPCR analysis. We found that SAG stimulation resulted in increased expression of *SHH* (*P* = .0156) and its downstream target *PTCH1* (*P* = .0312) compared to vehicle-treated controls (**Fig. 5o – u**), confirming activation of Hh signaling in degenerated human NP cells. We also found that activation of Hh signaling increased the expression of *ACAN* (*P* = .0312, **Fig. 5q**) with a corresponding significant decrease in expression of *MMP13* (*P* = .0156, **Fig. 5r**). Moreover, SAG treatment significantly inhibited the expression of neurotropic and inflammatory mediators including *NGF* (*P* = .0156, **Fig. 5s**), *TAC1* (*P* = .0312, **Fig. 5t**), and *IL1B* (*P* = .0156, **Fig. 5u**). These results indicate that SAG-mediated activation of Hh signaling can partially shift degenerated human NP explants toward a more anabolic and less inflammatory transcriptional profile.

We next tested whether the GSK3β inhibitor BIO, previously reported to activate Hh-associated signaling in mouse discs ^7^, could modulate degenerative transcriptional programs in human NP explants. NP tissue collected from each subject was split into two groups; one group was stimulated with BIO (10 µM), and the control group was vehicle treated. We found that stimulation of Grade III and IV NP explants with BIO resulted in a robust Hh signaling response by positively regulating the expression of its target *Ptch1* (*P =* .0078), and NP marker *TBXT* (*P =* .0156) compared to vehicle treated controls (**Extended data Fig. 6a** and **b**). Moreover, inhibition of GSK3beta stimulated mRNA expression of ECM marker *ACAN* (*P =* .0234) and suppressed the expression of matrix degrading enzymes like *MMP13* (*P =* .0078), and *ADAMTS4* (*P =* .0156, **Extended data Fig. 6c – e**). Furthermore, we also observed beneficial effects of stimulating Hh signaling via inhibition of GSK3beta by decreased expression of neurotropic and inflammatory mediators including *NGF* (*P* = .0156)*, TAC1* (*P* = .0312)*, IL1B* (*P =* .0078), and angiogenic factor *VEFGA* (*P* = .0391, **Extended data Fig. 6f – i**).

Together, these results indicate that Hh-associated signaling is pharmacologically modifiable in degenerated human NP explants, and supports its therapeutic relevance, although pathway specificity, durability and safety will require further validation before therapeutic translation.

## DISCUSSION

This study employed independent genetic approaches and lineage-tracing strategies to show that age-related pathological degeneration of the intervertebral disc results from the loss of *Shh*-expressing NP cells. Decline in SHH expression and NP cells is most dramatic in the lumbosacral disc between L5 to S1 levels. The persistence of NP-lineage cellular debris suggests that advanced aging impair clearance of NP remnants, potentially through reduced diffusion across the endplate, altered matrix turnover or impaired autophagy (reviewed by ^50^).

While the entire NP originates exclusively from the *Shh*-expressing notochord, we found that NP cells are molecularly heterogeneous as most NP cells turn off *Shh*-expression soon after birth. An *Shh*-expressing NP subset persists into aging until advanced NP-lineage depletion, suggesting that a specialized NP cell population may sustain paracrine Hh signaling in the adult disc. Loss of *Shh*-expressing NP cells was accompanied by reduced Hh target expression in both NP and AF compartments and by altered ECM turnover and increase in inflammatory and neurotrophic markers. Lineage tracing of sensory neurons further showed axonal sprouting and nerve fiber ingrowth into severely degenerated aged discs with reduced SHH expression.

Conditional *Shh* deletion in middle-aged mice demonstrated that loss of NP-derived SHH is sufficient to accelerate structural and molecular features that resemble natural disc aging. Conditional targeting of *Shh* resulted in an accelerated aging phenotype characterized by dramatic histopathological changes including changes in reticular-shaped NP into CLC syncytium by 2.5 months and loss by 5 months later. Because NP-specific *Shh* deletion reduced Hh target expression in both NP and AF cells, NP-derived SHH likely acts as a paracrine coordinator of multicompartment disc homeostasis. Moreover, genetic targeting of *Shh* resulted in NP cell death and increased expression of inflammatory markers and neurotropic factors by both NP and AF cells. A recent study showed a cell-autonomous role of SHH in NP cells ^48^, demonstrating that SHH directly regulates NP cells.

Furthermore, transient NP-specific *rShh* overexpression in middle-aged mice delayed age-associated degenerative remodeling of the lumbosacral disc. Compared with littermate controls, transient rSHH overexpression preserved NP morphology, ECM-associated markers and disc architecture while reducing molecular features of catabolic and inflammatory remodeling. These changes included higher expression of NP markers and ECM by both NP and AF cells, and decreased expression of catabolic markers including inflammatory and neurotropic factors by both NP and AF cells. These results indicate that enhancing SHH signaling in middle age can prevent the emergence of multiple age-associated degenerative features in mouse discs.

In addition, we found that human NP tissue showed similar reduction in SHH expression with age and increasing degeneration grade. As noted in our mouse studies, a decline in *SHH* expression in human NP cells was associated with declines in the expression of downstream *SHH* targets including *PTCH1*, *KRT19*, *TBXT*, and ECM markers. Reduced SHH expression with increased disc degeneration was also associated with increased inflammatory and neurotrophic mediators previously implicated in painful disc degeneration. From previous studies we know that increased expression of inflammatory molecules is related to disc degeneration in patients ^16–21^. Also, previous studies have reported increased innervation and vascularization of degenerated discs, which in a healthy state is an avascular and aneural tissue ^16–21^. Together, the mouse and human data support conservation of an age-associated decline in SHH pathway activity during disc degeneration. These changes were associated with a decline in the expression of the key developmental molecule, SHH, which is critical for the formation of the disc and its postnatal maintenance. Moreover, we showed that NP tissues from degenerated human disc could be stimulated and rejuvenated using small molecule activator of Hh signaling, and GSK3beta inhibitor that activates both Wnt and Hh signaling pathways. We also observed that inhibition of GSK3beta had a more robust effect than SAG in stimulating ECM related gene expression and inhibiting expression of inflammatory and neurotrophic factors which in line with previous observation that Wnt signaling is upstream of SHH signaling in mouse discs ^7^.

Overall, this study shows the important role of Hh signaling in regulating the homeostasis of the intervertebral disc and its rejuvenation. It is established that ECM, especially ACAN plays a critical role in preventing neovascularization ^51^ and innervation ^52^ of the disc. Here we show that SHH positively regulates ECM production by NP and AF cells and prevents expression of neovascularization and neurotropic factors. Hence, loss of SHH with pathological aging results in decreased ECM turnover and the disc becomes conducive to innervation and vascularization with corresponding higher levels of neovascularization and neurotropic factors. Also, considering the pathological disc express higher levels of inflammatory factors, and innervation of such disc it leads to pain pathology. In summary, our findings show that SHH, a key developmental molecule, is a direct regulator of disc homeostasis and is a potential target to treat the root cause of disc pathologies by healing and rejuvenating the disc tissue at molecular and structural levels.

## MATERIALS AND METHODS

### Mice

Male and female wild-type FVB and transgenic mice on FVB or mixed background were used. The following alleles were used in the current study: *Krt19t^m1(cre/ERT)Ggu^ (Krt19^CreERT/+^,^36,53^), (tetO)_7_CMV-rShh* (*rShh*,^54^), *Gt(ROSA)26Sor^tm3(ACTB-tdTomato,-EGFP)Luo^/J (R26^mT/mG^,^55^)*, *Shh^tm1(EGFP/Cre)Cjt/^J (Shh^Cre/+,56)^*, *Shh^tm2(cre/ERT2)cjt^/J (Shh^CreERT2/+,56^), Shh^tm2Amc^/J (Shh^flx/flx,57^), Shh-nLacZ^38^, Cg-E2f1^Tg(Wnt1-cre)2Sor^/J (Wnt1^Cre2/+,58^)*, and *Gt(ROSA)26Sor^tm1(rtTA,EGFPA)Nagy^* (*R26^rtTA/rtTA,59^*), *Gt(ROSA)26Sortm14(CAG-tdTomato)Hze/J* (*R26^TOM/+^*,^60^). Pups were genotyped within three weeks of birth by PCR using allele-specific primers and genomic DNA collected from tail biopsies. Tamoxifen (Sigma-Aldrich, USA, T5648) was prepared in warm corn oil (20 mg/mL) and dissolved by overnight incubation in a shaker at 37°C. Three doses of tamoxifen at 200 µg/g body weight were administered at 48-hour intervals to achieve efficient recombination and as previously described^36^. The mice were maintained in accordance with the National Institutes of Health Guide for the Care and Use of Laboratory Animals. All experiments adhered to institutional guidelines of the Institutional Animal Care and Use Committee (IACUC). Mice were maintained in a 12-hour day, 12-hour night cycle with food and water *ad libitum*.

#### Fate-mapping of neonatal mouse NP cells in aged mouse discs

To fate-map NP cells with age, postnatal (P) day five (P5) pups from an in-house generated *Krt19^CreERT/+^; R26^mT/mG^* line were subcutaneously injected with tamoxifen (200 µg/g body weight) using a 31-gauge syringe needle at P5, P7, and P9 as previously described^36^. Mice were euthanized at either P12 (*n* = 3) to validate efficient recombination or around 24 months (*n* = 3) of age to determine the fate of NP cells in aged mouse lumbar discs. Spines were collected at these time points and processed for cryosectioning. The *Krt19* allele was chosen to target the NP for two reasons: 1) *Krt19* is expressed explicitly by NP cells in the musculoskeletal system^36^, and 2) *Krt19^CreERT/+^* mediated recombination is very efficient in NP cells upon tamoxifen-induction^36^.

#### Fate-mapping of NP precursor cells in aged disc

An independent genetic strategy was employed to determine the fate of the entire population of NP cells by in-house crossing of *Shh^Cre/+^* and *R26^mT/mG^* alleles and allowing the mice to age. *Shh^Cre/+^-*driven recombination occurs in the node/notochord, the embryonic precursors of NP cells^30^. In our studies, all NP cells were fate-mapped using *R26^mT/mG^* as the conditional dual fluorescent reporter. Mice were euthanized at about 18 (*n* = 3) and 24 (*n* = 3) months of age, and the lumbar spines were collected and processed for cryosectioning.

#### Tracing of nerve fibers in aging mouse discs

*Wnt1^Cre2/+^; R26^mT/mG^* mice were generated in-house to track axonal sprouting and innervation of discs in aging mice using the membrane-bound GFP reporter. Mice were euthanized at P7 (*n* = 3) and about 26 (*n* = 3) months of age. Lumbar spines and lumbar dorsal root ganglions (DRG) were collected at these time points and processed for cryosectioning.

#### Identification of Shh-expressing NP cells at a given age

1. *Shh-nLacZ reporter allele and beta-Galactosidase Staining:* Lumbar spine from P10, 12-, and 24-month-old *Shh-^nLacZ^* mice (*n* = 3/age) were fixed in buffered 0.4% paraformaldehyde (PFA, Sigma-Aldrich, P6148-500G) and 0.2% glutaraldehyde (Fisher Scientific, 50-262-08) at 4°C for four hours on a shaker. The spines from adult mice were decalcified before cryosections were prepared in coronal plane. For X-gal staining, the cryosections were washed twice for 10 minutes each in wash buffer (2 mM MgCl_2_, Sigma-Aldrich, USA, M8266) and permeabilized for 20 minutes in 0.2% IGEPAL® CA-630 (Sigma-Aldrich, USA, I8896), 0.01% NaDeoxycholate (Sigma-Aldrich, USA, D6750), and 2 mM MgCl_2_. Sections were incubated in β-galactosidase staining solution [35 mM K_3_Fe(CN)_6_ (Sigma-Aldrich, USA, 244023), 35 mM K_4_Fe(CN)_6_ (Sigma-Aldrich, USA, P3289), 2 mM MgCl_2_, 0.01% NaDeoxycholate 0.02% IGEPAL® CA-630, 1 mg/mL of β-galactosidase (Sigma-Aldrich, USA, B4252), 0.5 M EGTA (Sigma-Aldrich, USA, E3889) in PBS] in a humidified chamber at 37°C for four hours. Next, the sections were washed twice in wash buffer for 10 minutes each, counterstained with Nuclear Fast Red (Sigma-Aldrich, USA, N3020) for 10 minutes, dehydrated in an increasing gradient of ethanol, cleared in xylene, and mounted using xylene-based mounting medium. Images were captured using a Nikon Eclipse microscope.
2. *Shh^CreERT2/+^ mediated conditional activation of fluorescent reporter allele. Shh^CreERT2/+^*were crossed with *R26^mT/mG^* conditional dual fluorescent reporter. Three doses of tamoxifen on alternate days were administered to each mouse at 12-, 18-, or 24- (*n* = 3/age) months of age as described above. The lumbar spines were collected 48 hours after the last dose of tamoxifen and processed for cryosectioning.

#### Conditional targeting of Shh in middle-aged mouse NP cells

1. *Krt19 induced conditional Shh knockdown:* Both *Krt19* and *Shh* are specifically expressed by NP cells in the musculoskeletal system. Therefore, a *Krt19^CreERT/+^*allele was crossed with a *Shh^flx/flx^* allele to generate *Krt19^CreERT/+^; Shh^flx/flx^* (*Shh*-cKO) and *Shh^flx/flx^* (WT control) littermates. Mice at about 10 months of age were administered tamoxifen (200 µg/g body weight) by three oral gavages at 48-hour intervals. One set of *Shh*-cKO (*n* = 6) and WT (*n* = 7) littermates was euthanized about 2.5 months post-tamoxifen treatment. The lumbar spine at L5-S1 levels was collected and fixed for histology. The NP and AF cells were separately microdissected from the proximal lumbar discs of the same mice and processed for RNA isolation^61^. Another set of *Shh*-cKO (*n* = 3) and WT (*n* = 3) littermates was aged for five months post-tamoxifen treatment before the lumbar spines were collected and processed for histological analysis.
2. *Fate-mapping of Shh-expressing NP cells following conditional targeting of Shh:* An orthogonal approach and genetic strategy was employed to target *Shh* in *Shh*-expressing NP cells while fate-mapping these cells. *Shh^CreERT2/+^* mice were crossed with *Shh^flx/flx^* and *R26^mT/mG^* to generate *Shh**<u>^flx/^</u>** ^CreERT2/+^; R26^mT/mG^* (*Shh*-cKO, *n* = 3/age) and *Shh^CreERT2/+^; R26^mT/mG^* (WT control, *n* = 3/age) littermates. Three doses of tamoxifen (200 µg/g body weight) were administered at 48- hour intervals by oral gavage to approximately 11-month-old mice. Mice were euthanized three or four months post-tamoxifen administration, and the lumbar spines were collected and processed for histological analysis.

#### Conditional and transient overexpression of rSHH in the NP cells

*Shh* is only expressed by NP cells in the disc and spine. Consequently, the *Krt19^CreERT/+^* allele was employed to induce conditional and transient overexpression of *rShh* in NP cells. Towards this, *Krt19^CreERT/+^, R26^rtTA/rtTA^* and *(tetO)_7_CMV-rShh* lines were crossed to generate *Krt19^CreERT/+^*; *R26^rtTA/rtTA^*; *(tetO)_7_CMV-rShh* (*rShh*-GOF, *n* = 4) and *R26^rtTA/rtTA^*; *(tetO)_7_CMV-rShh* (WT, *n* = 3) littermates. Tamoxifen was administrated (three doses of 200 µg/g body weight at 48-hour intervals) at about 12 months of age to induce the expression of the *R26^rtTA^* and GFP in the NP cells. Shortly after, the mice were administered doxycycline hyclate (Sigma-Aldrich, USA, D9891) at 200 µg/mL concentration in amber-colored drinking water bottles for one month. Upon administration of doxycycline, the *(tetO)_7_CMV-rShh* allele causes the conditional and transient overexpression of a rat *Shh* cDNA (rSHH) in the Cre-expressing driver cell population^54^. Fresh doxycycline water was replaced every two days. Tamoxifen- and doxycycline-treated WT littermates were used as controls. The mice were euthanized six months later when they reached about 19 months of age. At this time the lumbosacral levels (L5-S1) were collected, fixed, and processed for histology and entire discs from the proximal lumbar levels of the same mice were collected and processed for RNA isolation.

### Human NP tissue

#### Human NP tissue collection

NP tissue was collected using the hospital’s IRB-approved research study. Patients recruited for this study were undergoing spinal surgery due to a prior medical diagnosis and treatment. The patients were informed about the study and provided written consent for post-surgical collection of NP tissue, which otherwise would have been discarded after surgery. The recruited subjects included males and females from 19-79 years of age. A total of 36 samples were analyzed in the current study. The MRI images of the lumbar spine of each patient, which were collected as a part of the routine medical examination before surgery, were graded for disc degeneration by two blinded radiologists from the hospital who used the Pfirrmann grading system^49^. Following surgery, the NP tissue samples were immediately stored on ice and delivered to the lab to be weighed, washed three times in PBS, and processed for culture, RNA isolation, or fixed in buffered 4% PFA for cryosectioning followed by immunostaining.

#### Culture of human NP tissue

NP tissues collected from the lumbar discs with a Pfirrmann score of III or IV were subject to culture. About 50 mg of NP explant was cultured per well in 6-well culture plates using serum-free DMEM/F12 medium (Fisher Scientific, USA, 11-039-021) supplemented with insulin-transferrin-sodium selenite (ITS, Roche Diagnostics, 110744547001) and 1% pen/strep/nystatin. Cultures were maintained at 37°C with 5% CO_2_. The NP explant from each subject (*n* = 7) was split into two cohorts and treated with either 5 µM SAG (Smoothened Agonist, Millipore, 566660) or vehicle (control). In another set of experiments, NP explants (*n* = 8) from each subject were split into two cohorts and treated with either vehicle only or with 10 µM BIO (6-bromoindirubin-3”-oxime, Stemgent, 04-0003), a small molecule GSK3b inhibitor, which has been shown to activate hedgehog signaling downstream of the ligand in middle-aged mouse discs in vitro^7^. Fresh medium, along with supplements and treatments, were changed daily. Cultures were maintained for about 2.5 days. At the end of the culture experiment, the NP explant was washed three times in PBS and stored in RNA*later*^TM^ for downstream analysis.

### Histological analysis

#### Histology of mouse lumbar spine

The lumbar spine was dissected in cold PBS and lumbosacral levels were immediately fixed in buffered 4% PFA at 4°C for six hours. Following three washes in PBS at 4°C, the spines of skeletally mature mice were decalcified for seven to nine days in 0.5M ethylenediaminetetraacetic acid (EDTA, Sigma-Aldrich, USA, E9884), pH 7.4 at 8°C on a shaker. Spines were washed three times for 30 minutes each in cold PBS and molded in a coronal plane using Tissue-Tek® optimum cutting temperature (O.C.T, VWR, USA, 102094-106). Molds were snap-frozen and stored at -80°C until further use. A Leica cryostat was used to prepare 8 µm thick cryosections in the coronal plane, which were stored at -80°C until further use.

Mid-coronal sections were H&E stained using standard protocol to analyze histological changes. Two blinded raters scored histopathological changes in the NP, AF, EP, and interface regions using H&E-stained sections and a point-based mouse intervertebral disc histopathological scoring system^62^. The histopathological score of each disc was determined using the average score of two raters. Histopathological changes between cohorts for each disc compartment were analyzed using mixed-effect analysis followed by Šídák’s multiple comparisons test.

#### Histology of human NP tissue

NP tissue was washed in cold PBS and fixed in 4% PFA at 4°C on a shaker overnight. The following day, the samples were washed three times in cold PBS at 4°C, and molds were prepared using O.C.T. A Leica cryostat was used to prepare 8 µm thick cryosections, which were stored at -80°C until further use. Standard H&E staining was carried out to determine histological changes. Images were captured using a Nikon Eclipse wide-field microscope and NIS Elements AR software.

### Immunofluorescence and lineage-tracing analysis

#### Immunofluorescence analysis

Immunostaining was performed on cryosections prepared from mouse lumbar spine and human NP tissue. The slides were air-dried, washed twice in PBS, and permeabilized with 0.25 or 0.5% Triton X-100 prepared in PBS (PBST) at room temperature. Next, the sections were blocked in blocking buffer [10% Donkey serum (Jackson ImmunoResearch, USA, 017-000-121), 4% IgG-free BSA (Jackson ImmunoResearch, USA, 001-000-162), and 0.1% PBST] at room temperature for one hour in a humidified chamber. Next, the sections were hybridized with specific primary antibodies (Extended data Table 1) that were diluted in blocking buffer and incubated in a humidified chamber at 4°C overnight. Next day, the slides were washed in PBS three times for five minutes each. To visualize the protein of interest, sections were hybridized with Alexa Flour® conjugated secondary antibodies (Extended data Table 2, Jackson ImmunoResearch, USA) that were diluted in blocking buffer and incubated in a humidified chamber in the dark at room temperature for one hour. Each experiment included a negative control slide that was only incubated with the fluorescence-tagged secondary antibody, and not the primary antibody, and used to offset background fluorescence for quantification. The sections were counterstained with DAPI (1:5000, Life Technologies, USA, D1306) for five minutes, washed in PBS, and mounted using ProLong^TM^ Gold Antifade Mountant (Life Technologies, USA, P36934). A minimum of three serial sections were imaged and quantified per biological replicate.

#### Lineage tracing

Cryosections were washed twice in cold PBS and counterstained with DAPI for five minutes. The slides were mounted using ProLong^TM^ Diamond (Life Technologies, USA, P36962), and a minimum of six serial sections were imaged per biological replicate. Lineage-tracing by analyzing the GFP expression in the *R26^rtTA/rtTA^* allele was performed by immunostaining due to weak endogenous GFP expression.

#### RNAscope

RNAscope® ISH Fluorescent Assay v2 (Advanced Cell Diagnostics, Newark, CA, 323120) with high specificity and sensitivity was used to determine *Shh* mRNA expression in NP cells of mouse IVDs following the manufacturer’s instructions. Lumbar spines were dissected from P7 FVB mice (*n* = 3) and fixed in 4% PFA at 4°C for 24 hours. The spines were washed three times in PBS and processed for cryosectioning as described above. The slides were equilibrated to room temperature, postfixed with 4% PFA at 4°C for 15 minutes, and dehydrated in an ethanol series followed by target retrieval at 99°C for five minutes (in 1X target Retrieval reagent). Slides were pre-treated with H_2_O_2_ at room temperature for 10 minutes before protease III treatment at 40°C for 30 minutes. RNAscope® ISH Fluorescent Assay to detect *Shh* mRNA was performed using Mm-*Shh*-C2 probe (Advanced Cell Diagnostics, Newark, CA, 314361-C2) which was diluted with 100 µL of probe diluent, warmed at 40°C for 10 minutes, and applied to slides which were then incubated at 40°C for two hours in a humidity control tray (Advanced Cell Diagnostics, Newark, CA, PN 310012) placed in a HybEZ IITM hybridization system (Advanced Cell Diagnostics, Newark, CA, PN 321710). The slides were washed twice in 1X wash buffer, followed by hybridization at 40°C with amplifier AMP1 for 30 minutes, AMP2 for 30 minutes, and AMP3 for 15 minutes. Next, the slides were washed, and the signal was developed using Multiplex FL v2 HRP C2 at 40°C for 15 minutes and Atto TM 550 at 40°C for 30 minutes. The slides were then washed in 0.1M PBS (pH 7.4), nuclei were counterstained with DAPI (1:5000), and mounted using ProLong™ Gold Antifade Mountant (ThermoFischer Scientific, P36935). Images were captured at 60X magnification using a Ni Eclipse fluorescence microscope (Nikon, Tokyo, Japan) and NIS Elements AR software.

#### TUNEL assay

Cell death was determined using the *In Situ* Cell Death Detection Kit, TMR red (Millipore Sigma, USA, 12156792910) following the manufacturer’s protocol. The sections were counterstained with DAPI for five minutes and mounted using ProLong^TM^ Gold antifade moutant. A minimum of three serial sections were imaged and quantified per biological replicate.

#### Fluorescence microscopy

Fluorescence imaging was performed using a Zyla sCMOS monochromatic digital camera and DAPI, GFP, TxRD, and Cy5 filter cubes on a Nikon Eclipse Epi-fluorescence microscope and NIS Elements AR software (Nikon, Japan). Images were captured at 10x, 20x, and 60x magnification. The DIC filter was used for dark-field imaging to determine structural changes and to identify disc regions. High magnification (60x) images were deconvoluted by NIS Elements AR software using the Landweber algorithm and 20 iterations.

### Quantification of fluorescence data

Immunofluorescence data was quantified using Nikon NIS Elements AR software (Nikon, Japan).

#### Immunofluorescence intensity

Protein expression was measured by levels of immunofluorescence intensities for proteins localized in the extracellular space (SHH, ACAN, CSPG, COL1A1, COL2A1, and COLXA1). To quantify protein expression, the selected region of interest (ROI) was specific to an area that was ubiquitously occupied by NP or AF in Figures 2, 3, and 4. Mean fluorescence intensities (MFI) were quantified from entire disc for extracellular matrix-related markers in Figure 4, and Extended data in Figure 1. “ROI sum intensity” was divided by the “ROI area” to obtain the metric: total fluorescence intensity per μm^2^. Statistical analyses were performed using GraphPad Prism version 10. Data that compared three factors were analyzed using Brown-Forsythe and Welch ANOVA tests followed by Dunnett’s T3 multiple comparison tests. The differences between the two cohorts (age or experimental conditions) were analyzed using the unpaired *t-*test. *P* < .05 was considered statistically significant.

#### Quantification for number and percentage of NP cells

Changes with age among the total number of NP cells as well as the percentage of NP cells expressing a particular marker (in Figures 1, 2, 3, and 4, Extended Data Figure 1, 3) were quantified within an ROI encompassing the entire NP space. The total numbers of NP cells were determined by thresholding for DAPI for object count. Within that ROI, the numbers of NP cells expressing a specific marker (KRT19-IF+, Bra/T-IF+, TUNEL+, or GFP+) were counted by thresholding for the signal. These numbers were divided by the total number of NP nuclei to determine the percentages. Each experimental cohort used a minimum of three serial sections as technical replicates for each biological sample. Statistical analysis of both the percentages of NP cells for a specific marker and the changes in the total number of NP cells was performed using GraphPad Prism version 10. Further statistical analysis was conducted either using One-way ANOVA followed by Tukey’s multiple comparisons test, or Brown-Forsythe and Welch ANOVA followed by Dunnett’s T3 post-hoc analysis for multiple comparisons of three groups, or the unpaired *t-*test with Welch’s correction to compare two groups. *P* < .05 was considered statistically significant.

#### Quantification of disc vascularization

Vascularization of the intervertebral disc was quantified through the analysis of CD31 immunostaining and the NIS elements measuring tool. The distance and depth of CD31+ structures into the outer AF were divided by the total width of that disc. AF on both sides of each disc was quantified using three serial sections per biological replicate. Statistical analysis was performed using GraphPad Prism version 10 and the unpaired *t-*test. *P* < .05 was considered statistically significant.

### RNA isolation and qPCR analysis

For qPCR data presented in Figures 1, 2, and 3, the NP and AF components from the proximal lumbar discs of the mouse spine were microdissected in cold PBS under a Nikon stereomicroscope (SMZ1000, Nikon, Japan) and processed for RNA isolations using the published protocol^61^. For qPCR data presented in Figure 4, the entire disc from the proximal lumbar levels was collected. Samples were directly collected in RNA*later*^TM^ (Invitrogen by Thermo Fisher Scientific, Lithuania, AM7024) and stored at 4°C for 24 hours. For qPCR data of human NP tissue presented in Figure 5 and Extended data in Figure 6, the tissue was stabilized in RNA*later*^TM^ upon arrival in the lab or at the end of the culture experiment. Total RNA was isolated by extraction in TRI-reagent and followed by purification and elution using a Qiagen RNeasy RNA isolation kit. RNA concentration was quantified in duplicate using a NanoDrop™ One Microvolume UV-Vis Spectrophotometer (Thermo Scientific, USA, AZY1601393). RNA quality was assessed by Agilent 2100 Bioanalyzer (Agilent Technologies, Inc. Santa Clara, CA) at the Genomics Core facility of the institute. RNA was converted into cDNA using SuperScript™ IV First-Strand Synthesis System (Invitrogen by Thermo Fisher Scientific, Lithuania, 18091050). Multiplex qPCR was performed using the CFX96 Touch™ Real-Time PCR Detection System (Bio-Rad, Singapore, 1855195). Each PCR reaction used 8 ng of mouse cDNA or 16 ng of human cDNA, iQ™ Multiplex Powermix (Bio-Rad, USA, 1725849) master mix, gene- and species-specific TaqMan^TM^ probes (Extended data Table 3) conjugated to FAM-MGB, and primer-limited internal controls (Beta2 Microglobulin for mouse and GAPDH for human) conjugated to VIC-MGB (Extended data Table3). Data are presented as Mean ± S.D. and relative to the reference gene.

For analyzing changes in gene expression GraphPad Prism version 10 and Brown-Forsythe and Welch ANOVA tests followed by posthoc analysis using Dunnett’s test for multiple comparisons were employed for comparing expression at 12-, 18-, and 24-months of age in Figures 1 and 2, unpaired *t-*test between control and experimental cohorts in Figure 3, 4, and Figure 5h, Kruskal-Wallis test followed by Dunn’s multiple comparison tests in Figure 5i – n, and Wilcoxon matched-pairs signed rank test for qPCR data presented from NP explant culture experiment in Figure 5o – u and Extended data Figure 6a – i. The difference between gene expression in the vehicle- and SAG- (or BIO-) treated samples of the same biological replicate is indicated by a connecting line between the two cohorts in qPCR data presented in Figure 5o – u and Extended data Figure 6a – i. Changes in gene expression between two cohorts, when no amplification was detected in replicates of a given cohort (*Adamts5* and *Mmp13* in Figure 4o, *Bdnf* and *Nfg* in Figure 4p), were analyzed using one sample *t* and Wilcoxon test with a hypothetical value 1. *P* < .05 was considered statistically significant.

### Statistical Analysis

All statistical analyses were conducted in GraphPad Prism v10. Unless otherwise stated, *P* < .05 was considered statistically significant. Details for each analysis are specified in the respective methods section and figure legends. When multiple comparisons were conducted, the *P* values in the results section reflect the main ANOVA outcome.

## ACKNOWLEDGEMENTS

We sincerely thank Drs Alex Joyner (Memorial Sloan Kettering, New York, USA) and Chris Wylie (Cincinnati Children’s Hospital Medical Center, Cincinnati, USA) for insightful discussions during this study. We also sincerely thank Mr. Ravij Mehta for critical review of the manuscript. The TROMA-III antibody against CK19 developed by Kemler, R., was obtained from the Developmental Studies Hybridoma Bank, created by the NICHD of the NIH and maintained at The University of Iowa, Department of Biology, Iowa City, IA 52242.

## AUTHOR CONTRIBUTION

Conceptualization, C.L.D.; Methodology, C.L.D.; Validation, S.M., R.A., P.P., C.L.D.; Formal Analysis, C.L.D., E.B.; Investigation, S.M., R.A., P.P., R.P.., S.L., C.L.D.; Resources, C.L.D.; J.C.F., R.C.H., D.R.L., B.A.R., H.J.K., H.S.S., M.E.C., S.Q., T.J.A.; Data Curation, S.M., C.L.D.; Writing – Original Draft Preparation, S.M., C.L.D.; Writing – Review & Editing, all authors.; Visualization, S.M. and C.L.D.; Supervision, C.L.D.; Project Administration, C.L.D.; Funding Acquisition, C.L.D.; All authors read and approve of the manuscript.

