## Extended Table 1 and 2 for "Sonic Hedgehog Is An Important Regulator Of Intervertebral Disc Homeostasis And Rejuvenation"

**Extended data Table 1.** Primary antibodies used for immunofluorescence analysis**.**

| **Description of antigen** | **Gene ID** | **Isotype** | **Vendor** | **Catalog #** | **Dilution** | **Concentration** |
| --- | --- | --- | --- | --- | --- | --- |
| Keratin 19, cytokeratin 19 | KRT19 | Rat IgG2a | DSHB Hybridoma | TROMA-III, RRID:AB_2133570 | 1 in 100 | 0.257 mg/ml |
| Sonic hedgehog | SHH | Rat IgG2a | Sigma- Aldrich | S4944 | 1 in 50 | 0.5 mg/ml |
| Brachyury | T, TBXT, BRA | RP IgG | Abcam | ab20680 | 1 in 10 | 1.0 mg/ml |
| Collagen Type I Alpha 1 Chain | COL1A1 | RP IgG | GeneTex | GTX41285 | 1 in 100 | 1.0 mg/ml |
| Collagen Type II Alpha 1 Chain | COL2A1 | MM IgG2a | Novus Biologicals | NB600-844 | 1 in 100 | 0.2 mg/ml |
| Collagen Type X Alpha 1 Chain | COLXA1 | RP IgG | Millipore Sigma | 234196 | 1 in 100 | Undiluted serum |
| Chondroitin sulfate | CSPG, CHSO4 | MM IgM | Abcam | ab11570 | 1 in 100 | 1.6 mg/ml |
| Aggrecan | ACAN | MM IgG1 | Invitrogen | AHP0022 | 1 in 100 | 1 mg/ml |
| Platelet And Endothelial Cell Adhesion Molecule 1 | CD31, PECAM1 | Goat IgG | R&D | AF3628 | 1 in 50 | 0.2 mg/ml |
| GFP | GFP | RP IgG | Life Technologies | 11122 | 1 in 100 | 2 mg/mL |

**Extended data Table 2.** Fluorophore-tagged secondary antibodies.

| **Secondary Antibodies** | **Catalog #** | **Dilution** | **Concentration** |
| --- | --- | --- | --- |
| Alexa Fluor® 594-AffiniPure Goat Anti-Rabbit IgG (H+L) | 111-585-144 | 1 in 200 | 1.5 mg/ml |
| Alexa Fluor© 647-AffiniPure donkey anti-rabbit IgG (H+L) | 711-605-152^a^ | 1 in 200 | 1.5 mg/ml |
| Alexa Fluor® 647-AffiniPure Goat Anti-Mouse IgG (H+L) | 115-605-146^a^ | 1 in 200 | 1.5 mg/ml |
| Alexa Fluor® 647-AffiniPure Goat Anti-Rat IgG (H+L) | 112-605-062^a^ | 1 in 200 | 1.5 mg/ml |
| Alexa Fluor© 647-AffiniPure Donkey Anti-Mouse IgM, μ Chain Specific | 715-605-020^a^ | 1 in 200 | 1.5 mg/ml |
| Alexa Fluor® 488-AffiniPure Donkey Anti-Goat IgG (H+L) | 705-545-147^a^ | 1 in 200 | 1.5 mg/ml |
| Alexa Fluor® 488-AffiniPure Donkey Anti-Mouse IgM, μ Chain Specific | 715-545-140^a^ | 1 in 200 | 1.5 mg/ml |

Footnote: ^a^ Vendor, Jackson ImmunoResearch Laboratories, USA.
