## Extended Table 3 for "Sonic Hedgehog Is An Important Regulator Of Intervertebral Disc Homeostasis And Rejuvenation"

| **Extended data Table 3.** List TaqMan^TM^ probes used for gene expression analysis (Thermo Fisher Scientific, USA) | | | |
| --- | --- | --- | --- |
| **Description** | **Gene Symbol/ ID** | **TaqMan probe ID:** | **Dye** |
| **MOUSE** | | | |
| Sonic hedgehog | *Shh* | Mm00436528_m1 | FAM-MGB |
| Protein patched homolog 1 | *Ptch1* | Mm00436026_m1 | FAM-MGB |
| Glioma-Associated Oncogene Homolog 1 | *Gli1* | Mm00494654_m1 | FAM-MGB |
| Keratin 19, Cytokeratin 19 | *Krt19* | Mm00492980_m1 | FAM-MGB |
| Brachyury | *T* | Mm01318252_m1 | FAM-MGB |
| SRY-Box Transcription Factor 9 | *Sox9* | Mm00448840_m1 | FAM-MGB |
| Aggrecan | *Acan* | Mm00545794_m1 | FAM-MGB |
| Collagen Type I Alpha 1 Chain | *Col1a1* | Mm00801666_g1 | FAM-MGB |
| Collagen Type II Alpha 1 Chain | *Col2a1* | Mm01309565_m1 | FAM-MGB |
| ADAM Metallopeptidase With Thrombospondin Type 1 Motif 4 | *Adamts4* | Mm00556068_m1 | FAM-MGB |
| ADAM Metallopeptidase With Thrombospondin Type 1 Motif 5 | *Adamts5* | MM00478620_m1 | FAM-MGB |
| Matrix Metallopeptidase 13 | *Mmp13* | MM00439491_m1 | FAM-MGB |
| Brain Derived Neurotrophic Factor | *Bdnf* | Mm04230607_s1 | FAM-MGB |
| Nerve Growth Factor | *Ngf* | Mm00443039_m1 | FAM-MGB |
| Tachykinin Precursor 1 | *Tac2, Tac1* | Mm01160362_m1 | FAM-MGB |
| Vascular Endothelial Growth Factor A | *Vegfa* | Mm00437306_m1 | FAM-MGB |
| Interleukin 1 Beta | *Il1b* | Mm00434228_m1 | FAM-MGB |
| Tumor Necrosis Factor | *Tnf* | Mm00443258_m1 | FAM-MGB |
| Prostaglandin-Endoperoxide Synthase 2 | *COX2* | Mm03294838_g1 | FAM-MGB |
| Beta-2-Microglobulin (*reference gene*) | *B2m* | Mm00437762_m1 | VIC-MGB |
| **HUMAN** | | | |
| Sonic hedgehog | *SHH* | Hs00179843_m1 | FAM-MGB |
| Protein patched homolog 1 | *PTCH1* | Hs00181117_m1 | FAM-MGB |
| Keratin 19 | *KRT19* | Hs00761767_s1 | FAM-MGB |
| Brachyury | *TBXT* | Hs01084475_g1 | FAM-MGB |
| SRY-Box Transcription Factor 9 | *SOX9* | Hs00165814_m1 | FAM-MGB |
| Aggrecan | *ACAN* | Hs00153936_m1 | FAM-MGB |
| ADAM Metallopeptidase With Thrombospondin Type 1 Motif 4 | *ADAMTS4* | Hs00192708_m1 | FAM-MGB |
| Matrix Metallopeptidase 13 | *MMP13* | Hs00942584_m1 | FAM-MGB |
| Brain Derived Neurotrophic Factor | *BDNF* | Hs00380947_m1 | FAM-MGB |
| Nerve Growth Factor | *NGF* | Hs00171458_m1 | FAM-MGB |
| Tachykinin Precursor 1 | *TAC1* | Hs00243225_m1 | FAM-MGB |
| Vascular Endothelial Growth Factor A | *VEGFA* | Hs00900055_m1 | FAM-MGB |
| Interleukin 1 Beta | *IL1B* | Hs01555410_m1 | FAM-MGB |
| Glyceraldehyde-3-Phosphate Dehydrogenase (reference gene) | *GAPDH* | HS00266705_G1 | VIC-MGB |
