## Extended Data Figure 1 - 6 for "Sonic Hedgehog Is An Important Regulator Of Intervertebral Disc Homeostasis And Rejuvenation"

Extended Data Fig. 1

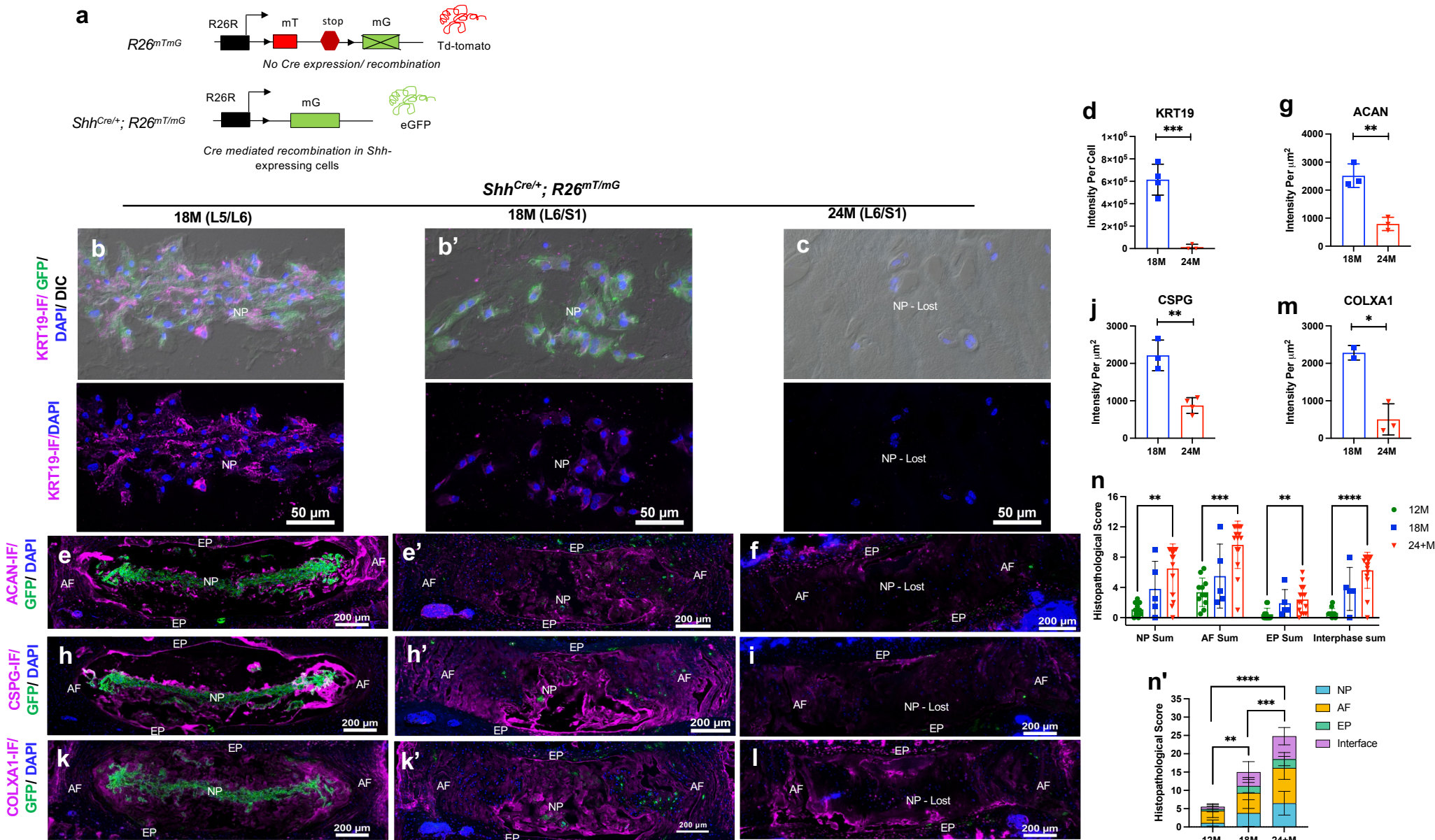

Extended Data Fig. 2

12M (L6/S1) *Shh<sup>nLacZ/+</sup>*

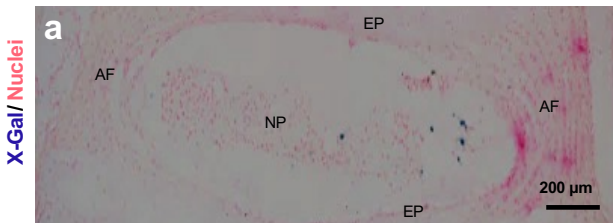

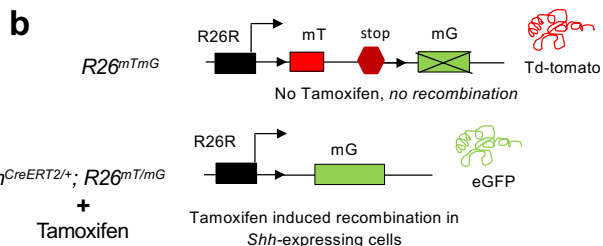

12M (L6/S1) *Shh<sup>CreERT2/+</sup>; R26<sup>mTmG</sup>*, Tam @ 12M

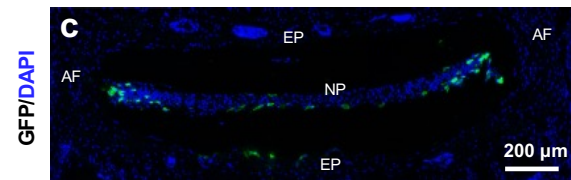

18M *Shh<sup>CreERT2/+</sup>; R26<sup>mTmG</sup>*, Tam @ 18M

L5/L6

L6/S1

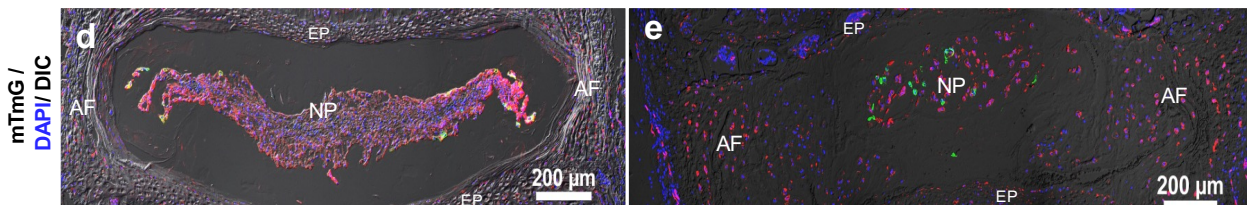

24M (L6/S1) *Shh<sup>CreERT2/+</sup>; R26<sup>mTmG</sup>* (Tam @ 24M)

L6/S1

L6/S1

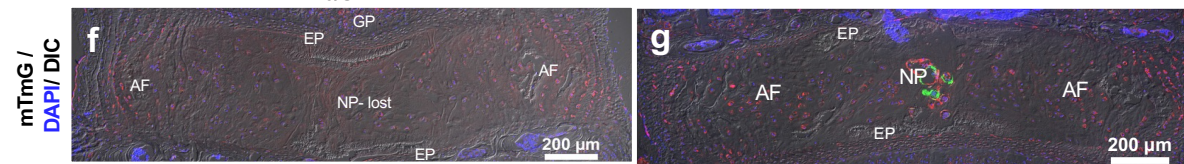

24M (L6/S1) *Shh<sup>CreERT2/+</sup>; R26<sup>mTmG</sup>*, Tam @ 24M

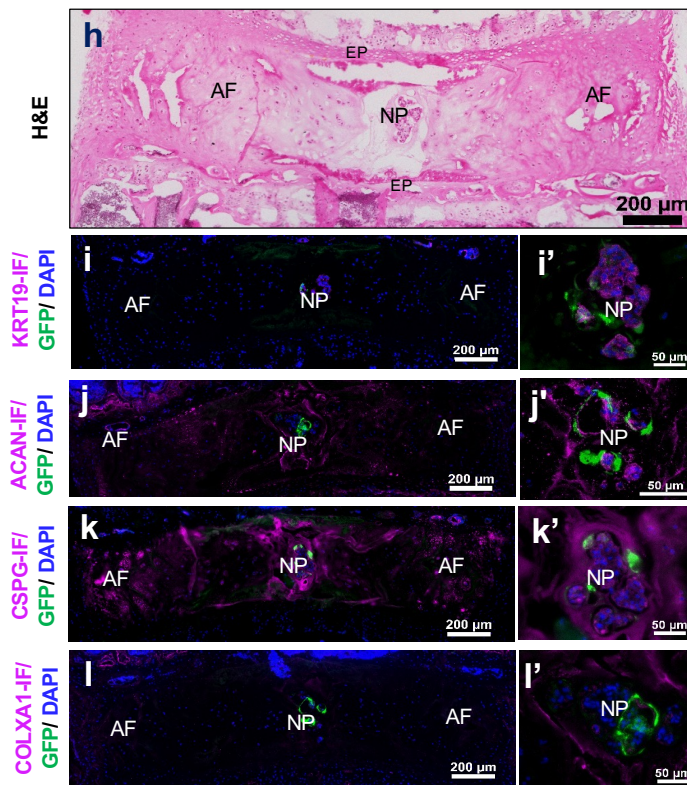

Extended data Fig. 3

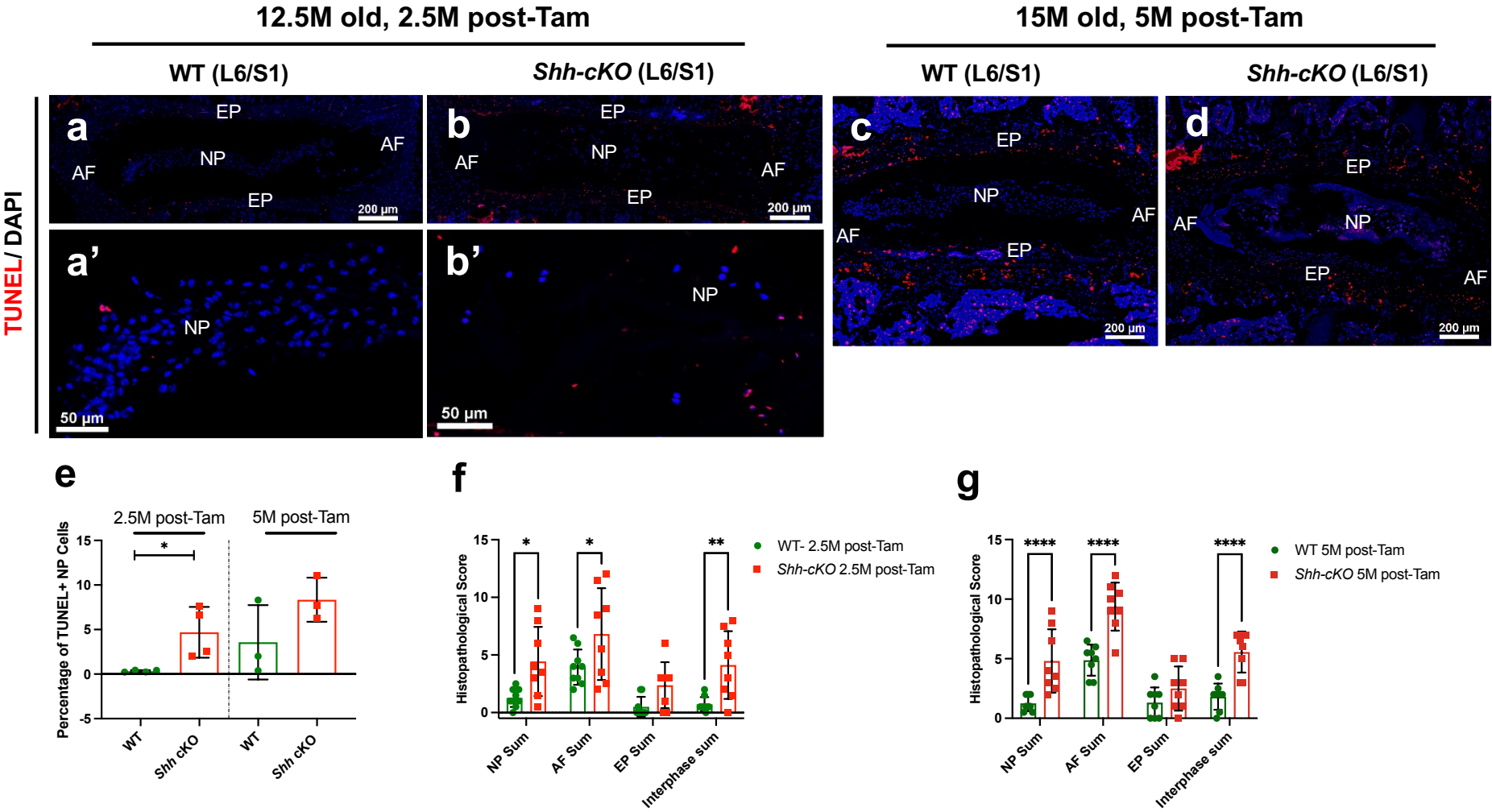

### Extended data Fig. 4

14M old, 3M post-Tam

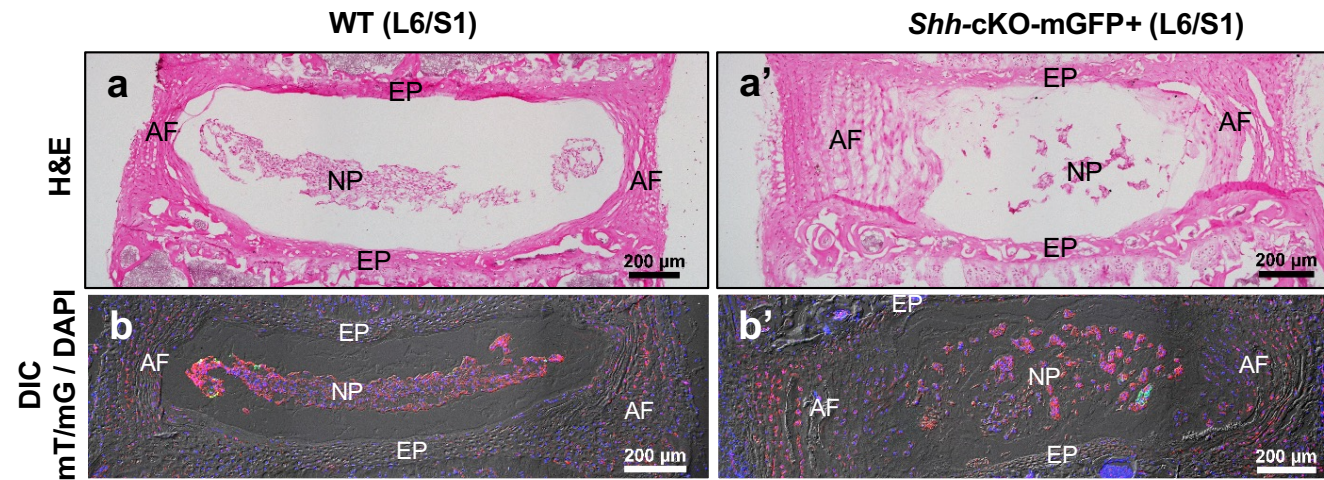

15M old, 4M post-Tam

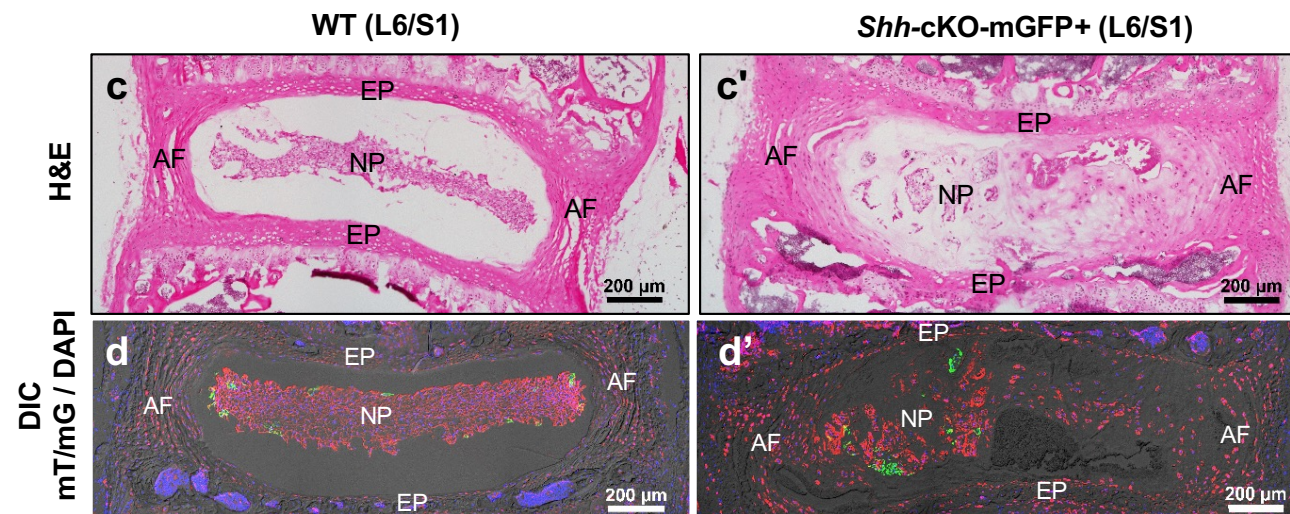

Extended data Fig. 5

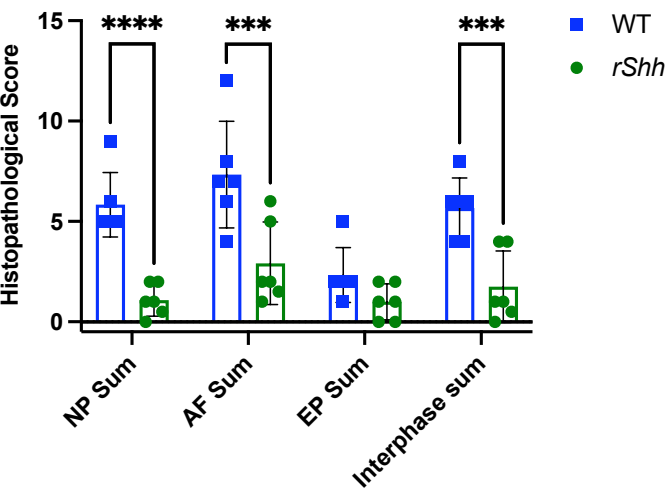

Extended data Fig. 6

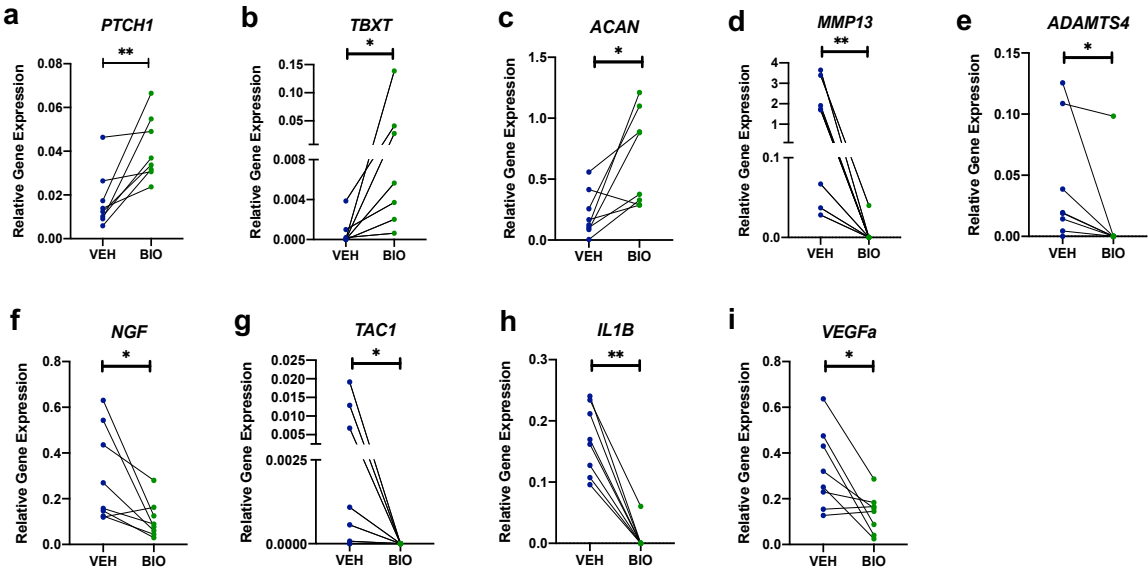

### FIGURE LEGENDS FOR EXTENDED DATA

**Extended Data Figure 1. Disc pathologies are associated with the loss of NP cells.** Schema of genetic recombination strategy to track node/notochord cells that give rise to NP cells (mGFP+) in the aged mouse lumbar discs (**a**). Mid-coronal section of lumbosacral discs of *Shh<sup>Cre/+</sup>; R26<sup>mT/mG</sup>* mice at 18M (**b, b'**) of age where all NP cells are mGFP+, and 24M (**c**) of age where mGFP+ NP cells are lost. Representative images showing immunostaining (purple) for KRT19 (**b, b', c**), ACAN (**e, e', f**), CSPG (**h, h', i**), and COLXA1 (**k, k', l**) with nuclei counterstained with DAPI (blue). Quantification of fluorescence intensity per cell for KRT19 (**d**) and immunofluorescence intensity for ACAN (**g**), CSPG (**j**), and COLXA1 (**m**) in the discs at 18M and 24M of age. Histopathological scores of lumbar discs from 12M, 18M and 24M old mice from various lineage-traced genetic cohorts presented in Figure 1, Figure 2, Extended Data Figure 1, and Extended Data Figure 2 showing the sum of each disc compartment (**n**) and cumulative score for the entire disc (**n'**) at a given age. Each dot represents a biological replicate in **d, g, j, m** and **n**. Statistical analysis by unpaired *t*-test. *n* = 3 mice per cohort (**d, g – m**), mixed-effects model (**n** and **n'**). Data are presented as mean ± S.D. \* *P* < .05; \*\* *P* < .01; \*\*\* *P* < .001; \*\*\*\* *P* < .0001. NP, nucleus pulposus. AF, annulus fibrosus. EP, end plate.

### **Extended Data Figure 2. Disc pathologies are associated with the loss of *Shh*-expressing NP cells.**

Representative image of a LacZ stained L6/S1 disc of 12M old *Shh<sup>nLacZ</sup>* reporter mice showing *Shh*-expressing (blue) NP cells in mid-coronal plane (**a**). Nuclei is counterstained with Nuclear Fast Red. Schema showing tamoxifen-inducible genetic recombination strategy to mark *Shh*-expressing NP cells at a given age in postnatal mouse disc (**b**). Mid-coronal section of the lumbosacral disc from the levels, age, and genotype indicated above each panel (**c – l'**). *Shh*-expressing cells (GFP+) at a given age were detected 48 hours following last tamoxifen treatment of *Shh<sup>CreERT2/+</sup>; R26<sup>mT/mG</sup>* mice at 12M (**c**), 18M (**d** and **e**), and 24M (**f – l'**) of age; nuclei counterstained with DAPI (blue). H&E-stained mid-coronal section of L6/S1 disc from a 24M old *Shh<sup>CreERT2/+</sup>; R26<sup>mT/mG</sup>* (shown in **g**). Immunostaining (purple) with KRT19 (**i** and **i'**), ACAN (**j** and **j'**), CSPG (**k** and **k'**), and COLXA1 (**l** and **l'**); nuclei counterstained with DAPI (blue). Images were captured at 20x magnification (**a, c, d, e, f, g, h, i, j, k, l**) and 60x magnification (**i', j', k', l'**). NP, nucleus pulposus. AF, annulus fibrosus. EP, end plate.

### **Extended Data Figure 3. SHH is crucial for the survival of NP cells.**

Mid-coronal section of lumbosacral (L6/S1) discs of WT (*Shh<sup>flx/flx</sup>*) and *Shh*-cKO (*Krt19<sup>CreERT/+</sup>; Shh<sup>flx/flx</sup>*) littermates at 12.5M of age, 2.5M post-tamoxifen induction (**a, a', b, b'**) and 15M of age, 5M post-tamoxifen induction (**c, d**). TUNEL (red, **a – d**) staining to determine cell death and nuclei counterstained blue with DAPI. Quantification of percentage of TUNEL+ NP cells in WT and *Shh*-cKO cohorts at 2.5M ($n = 4/\text{cohort}$) and 5M post-tamoxifen ($n = 3/\text{cohort}$) induction, with each dot representing a biological replicate. Quantification of histopathological scores of individual lumbosacral disc compartments of two discs per mouse for the 2.5M post-Tamoxifen (**f**) and 5M post-Tamoxifen (**g**) cohorts. Each dot represents a lumbar disc in **f - g**. Statistical analysis was conducted using an unpaired *t*-test (**e**) and mixed-effects model (**f** and **g**). Data presented as mean ± S.D. \* $P < .05$; \*\* $P < .01$; \*\*\*\* $P < .0001$. NP, nucleus pulposus. AF, annulus fibrosus. EP, end plate.

**Extended Data Figure 4. Fate-mapping of *Shh*-expressing NP cells following conditional targeting of *Shh* reveals differentiation of NP cells into multi-nucleated lacunae.**

Mid-coronal section of lumbosacral (L6/S1) discs of WT (*Shh<sup>CreERT2/+</sup>; R26<sup>mT/mG</sup>*) and *Shh*-cKO (*Shh<sup>flx/CreERT2/+</sup>; R26<sup>mT/mG</sup>*) littermates at 14M of age (3M post-tamoxifen induction, **a - b'**) and 15M of age (4M post-tamoxifen induction, **c - d'**). H&E staining of mid-coronal section (**a, a', c, c'**). Epifluorescence images captured with a dark field for fate-mapping mGFP+ *Shh*-expressing NP cells and analysis of changes in cell phenotype (**b, b', d, d'**). Nuclei counterstained blue with DAPI (**b, b', d, d'**). $n = 3$ mice per cohort per age. NP, nucleus pulposus. AF, annulus fibrosus. EP, end plate.

**Extended data Figure 5. Transient *rShh* overexpression prevents disc pathology.** The graph shows the histopathological score of each disc compartment. Two discs were scored per mouse and are represented by a dot on the graph. Statistical analysis by mixed-effect model. Data presented as mean ± S.D. \*\*\* $P < .001$; \*\*\*\* $P < .0001$.

**Extended Data Figure 6. GSK3β inhibitor activates hedgehog signaling and rescues the degenerative phenotype of human NP explants.** NP explants from human discs with Pfirrmann score III and IV were cultured for 2.5 days either in vehicle only (VEH) or stimulated with 10 μM BIO. Multiplex qPCR analysis for expression of indicated genes (PTCH1, **a**; TBXT, **b**; ACAN, **c**; MMP13, **d**; ADAMTS4, **e**; NGF, **f**; TAC1, **g**; IL1B, **h**; VEGFa, **i**) using *GAPDH* as an internal control. NP tissue from each subject was cultured under two conditions and is connected by a line to show pairs. Each dot represents a biological replicate. $n = 8$. Statistical analysis using Wilcoxon matched-pairs signed rank test (**a – i**). \* $P < .05$; \*\* $P < .01$.
